# Phageome transfer from gut to circulation and its regulation by human immunity

**DOI:** 10.1101/2024.06.05.597592

**Authors:** Aleksander Szymczak, Katarzyna Gembara, Stanisław Ferenc, Joanna Majewska, Paulina Miernikiewicz, Marek Harhala, Izabela Rybicka, Dominik Strapagiel, Marcin Słomka, Jakub Lach, Jan Gnus, Paweł Stączek, Wojciech Witkiewicz, Matlock Jeffries, Krystyna Dąbrowska

## Abstract

Bacteriophages dominate the human gut virome, yet their presence in the bloodstream remains orders of magnitude lower, suggesting that systemic dissemination is tightly regulated. How gut phages cross the intestinal barrier and which factors govern their persistence in circulation remain poorly understood, largely because prior studies characterized gut and blood viromes independently rather than in matched samples from the same individuals. Here we investigated phage translocation by shotgun metagenomics of matched colon mucosal biopsies and sera from 37 individuals with phage specific IgG profiling using a pan-phage proteome-derived phage display epitope library, complemented by an oral T4 model in mice. We found that phage abundance decreased by approximately 98% from intestinal mucosa to serum. The mucosal virome was dominated by Microviridae, which also accounted for most of the translocated phages. In the mouse model, phage titers dropped stepwise by ∼10^6^ fold from gut content to blood, with the sharpest reduction occurring at the mucosal-lymphatic interface. Among translocated viral operational taxonomic units 93.1% lacked taxonomic assignment, yet network analysis revealed reproducible co-enrichement with annotated families including Herelleviridae and Straboviridae, which showed significantly higher gut abundance among translocated observations (FDR <0.01). IgG reactivity against a specific phage in 90% of investigated individuals was associated with the absence of that phage in the patient’s virome; at the collective population analysis, IgG reactivity showed a weak negative association with serum phage abundance. These observations suggest antibody-mediated clearance that limits systemic persistence. Together, these findings suggest that rare epithelial passage, lymphatic trafficking, and IgG-mediated neutralization act as sequential filters that limit which gut phages reach and persist in the circulation, with implications for phage therapy delivery and for the dissemination of accessory genetic elements beyond the intestine.

## Introduction

Bacteriophages - viruses that infect bacteria - dominate the human intestinal virome reaching densities of 10^8^ - 10^10^ virions per gram of fecal content (Guerin and Hill 2020; Sutton and Hill 2019; Zuppi et al. 2021). Gut phages, called gut phageome, interact closely with bacteria to control their populations. Through predation and lysogeny, these viruses shape bacterial community structure, drive diversification and sustain niche competition that stabilizes microbial resilience against dysbiosis (Marantos, Mitarai, and Sneppen 2022; Easwaran et al. 2025). Furthermore, the gut phageome is dominated by temperate phages maintained as prophages, whose induction upon bacterial stress provides a constitutive supply of free virions available for dissemination (Shkoporov and Hill 2019; Dahlman et al. 2025).

The human phageome extends beyond the gastrointestinal tract. Phages have been detected across multiple tissues and organs, and their systemic presence is considered to depend on their ability to translocate into the bloodstream (Dąbrowska 2019; Bichet et al. 2021; Nguyen et al. 2017). Systemically dispersed phages can be linked to their bacterial hosts as shown in septic patients, where blood phages displayed pathogen-specific patterns (Bichet et al. 2021; Haddock, Barkal, and Bollyky 2023). Phage detection in blood is not limited to pathological conditions; phages can also be found in healthy individuals without active infections. However, the circulating phageome is orders of magnitude smaller than the gut reservoir and only trace levels of phage DNA are detected in blood (Nguyen et al. 2017; Moustafa et al. 2017; Majewska et al. 2019). While bacteriophages have been proposed to concentrate at mucosal surfaces through adhesion to mucin glycans (Barr et al. 2013; Almeida et al. 2019), this hypothesis is primarily supported by in vitro experiments and fish surface mucus models. Proposed routes of phage translocation to blood include transcytosis across epithelial barriers and vesicular transport. Active transcytosis of tailed phages across epithelial and endothelial monolayers has been clearly demonstrated in vitro and is considered a key mechanism for phage dissemination in the body (Bichet et al. 2021; Nguyen et al. 2017).

Bioavailability of phages disseminated in the body is determined by immune responses, involving both innate and adaptive immunity (Dąbrowska and Abedon 2019). Indeed, functional phage-neutralizing activity has been detected in sera of healthy individuals never exposed to phage therapy, with prevalence ranging from 11% for some Pseudomonas phages to 82% for coliphage T4 (Dąbrowska et al. 2014; Hodyra-Stefaniak et al. 2020; Kaźmierczak et al. 2021) suggesting routine systemic exposure to environmental and commensal phages. Conversely, the gut immune system maintains immunological tolerance toward commensal microorganisms, thereby sustaining microbial diversity and ecological balance (Belkaid and Hand 2014; Round and Mazmanian 2009). It remains unclear whether translocating bacteriophages induce specific immune responses or if they benefit from the phenomenon of tolerance. In particular, the ability of phages to trigger specific immune responses that may limit their translocation and systemic spread, as well as their ability to persist in the gut, remains largely unexplored.

A key limitation in our understanding of this problem is the lack of paired intestinal and systemic samples from the same individuals. Studies have characterized gut phageomes or circulating viromes separately, but without matched compartments it is difficult to determine which phages cross epithelial barriers and what are possible indicators of efficient or inefficient translocation. This challenge is compounded by viral dark matter up to 90% of virome sequences lack database homologs and the absence of a universal phage marker gene, making taxonomy-resolved tracking across body sites intractable with standard approaches (Zuppi et al. 2021; Badillo-Pazmay et al. 2025; Santiago-Rodriguez and Hollister 2022) Therefore, it is even more difficult to conclude how antibody responses relate to systemic detection of gut phages.

Here we analyze phage translocation from gut to circulation using matched mucosal biopsy and serum pairs from 37 individuals, combining metagenomic profiling with serological detection of phage-specific immunoglobulin G (IgG). We aimed to determine how different phage groups vary in their ability to translocate from the gut to the circulation. These observations stem from an animal model used to identify barriers that strongly affect phage translocation efficiency, leading to striking differences between transcytosis observed in vitro and the amounts of circulating phages or phage DNA detected in living individuals. We further assessed how specific recognition of phage antigens by serum antibodies is associated with phage presence in the gut and in circulation.

## Materials and Methods

### Study design and participants

Matched colon mucosal biopsies and peripheral blood were collected from adult patients undergoing diagnostic colonoscopy at the Endoscopy Laboratory of the Regional Specialist Hospital in Wrocław, Poland (Fig. 1). Participants were recruited based on gastrointestinal symptoms prompting colonoscopy. Individuals reporting antibiotic treatment within the preceding 6 months or diabetes treated with metformin were excluded. Intestinal biopsies were collected from regions showing no abnormalities on endoscopic examination. The cohort comprised 37 participants (21 women, 16 men; mean age 61 years; Supplementary Figure S1). The study was conducted in accordance with the Declaration of Helsinki and approved by the local Bioethical Commission of the Regional Specialist Hospital in Wrocław (KB/nr8/rok2017). Written informed consent was obtained from each participant.

**Figure 1.**
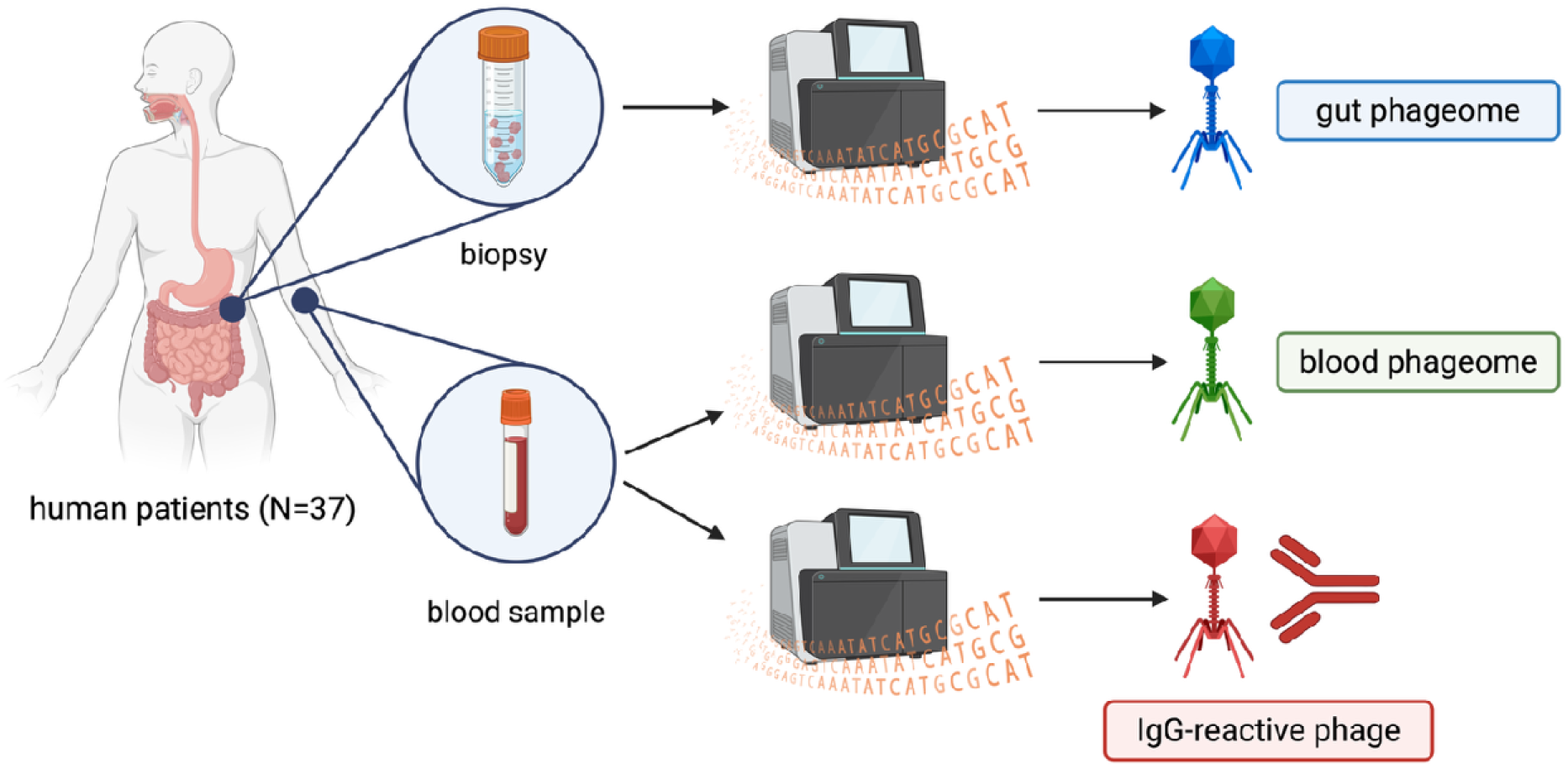
Human samples used in this study to investigate composition of gut and blood phageomes, and IgG antibody responses to the phageomes (BioRender).

### VLP enrichment and DNA preparation

Biopsies were incubated in PBS for 2 h and filtered through 0.22 μm membrane filters. Viral particles were enriched by CsCl density-gradient ultracentrifugation (step gradient: 1.7, 1.5, 1.35, and 1.15 g ml ¹) performed overnight at 62,000 × g at 4 °C in a swinging-bucket rotor. A 0.5 ml fraction between 1.35 and 1.5 g ml ¹ was collected by syringe for DNAse I treatment (A&A Biotechnology). Phage DNA isolation (Sherlock AX kit, A&A Biotechnology), was followed by whole-genome amplification (GenomiPhi V2, Cytiva). Whole blood was centrifuged at 2,000 RCF for 10 min at 4 °C. The serum layer was collected for total DNA isolation (Sherlock AX kit, A&A Biotechnology), followed by whole-genome amplification (GenomiPhi V2, Cytiva) before sequencing library preparation.

### Library preparation and sequencing

Shotgun libraries were prepared using a Vazyme DNA library preparation kit and sequenced as paired-end 2×150 bp on Illumina NextSeq 500/550 using the v2.5 Kit (Medium) at two facilities. Thirty-seven matched intestine-serum pairs were analyzed by shotgun sequencing. QC details after sequencing for samples available in Supplementary file - Sequences Quality Control.

### Read processing and host read removal

FASTQ files were quality-filtered and adapter-trimmed using fastp v0.24.2. Host reads were removed by alignment to human GRCh38.1 (UCSC) using Bowtie2 v2.4.5 using “-very-sensitive”; unmapped reads were retained.

### Assembly, vOTU definition, and abundance estimation

Non-host reads were assembled using metaSPAdes v4.20. Contigs <1 kb were excluded. Viral contigs were identified using VirSorter2 v2.2.4. vOTUs were defined by clustering contigs using CheckV v1.0.3 average nucleotide identity (ANI)-based clustering at 95% with 85% (AF) alignment fraction; representative sequences were selected per cluster prioritizing higher CheckV completeness, with ties broken by longer length. Reads were mapped to vOTU representatives with Bowtie2 (--very-sensitive). Alignments were filtered at mapping quality ≥30. Per-vOTU counts were converted to TPM: TPM = ((count /length)/Σ (count /length))×10. A vOTU was considered detected in a compartment only if breadth of genome coverage was ≥10%, and this detection filter was applied before defining within-subject status. For subject-level comparisons, vOTU abundance was summed separately across all intestine-derived samples and all serum-derived samples per subject. vOTUs were labeled “translocated” when summed intestine abundance >0 and summed serum abundance >0 within the same subject, and “only gut presence” when summed intestine abundance >0 and summed serum abundance =0.

To assess whether serum detection primarily reflected matched-pair biology rather than sporadic contamination, we quantified serum-intestine concordance and tested it against a pairing null. Of 2,488 vOTUs detected in serum across the cohort, 2,436 (97.9%) were also detected in at least one matched intestine sample, whereas 52 (2.1%) were detected only in serum. Moreover, within-subject serum-intestine overlap exceeded a permuted pairing null (mean overlap proportion 0.464 observed versus 0.405 expected; permutation P < 0.001; Supplementary Table S5), supporting that serum detections are non-random with respect to subject pairing.

### Taxonomy, host context, and lifestyle prediction

Taxonomic assignment was performed using against UHGV which contains collections from multiple databases (Metagenomic Gut Virus Compedium, Gut Phage Database, Metagenomic Mobile Genetic Elements Database, IMG Virus Resource v4, Hadza Hunter Gatherer Phage Catalog, Cenote Human Virome Database, Human Virome Database, Gut Virome Database, Atlas of Infant Gut DNA Virus Diversity, Circular Gut Phages from NCBI, Danish Enteric Virome Catalogue, Stability of the human gut virome and effect of gluten-free diet) with assignments requiring 95% ANI and 85% AF. Predicted lifestyle was obtained using BACPHLIP v0.96 with integrase annotations from PHROG and prophage predictions from geNomad v1.11.0.

### Gene prediction and functional annotation

Protein-coding genes were predicted using Prodigal v2.6.3. Functional annotation was performed with eggNOG-mapper v2.11.2 (Cantalapiedra et al., 2021) and, in parallel, phage-focused annotation using Pharokka v1.7.4 with PHROGs. KO/pathway summaries used mean values across CDS mapped to the same KO/pathway. Defense system calls used DefenseFinder v2.0.1.

### IgG-reactivity library and readout

A phage-derived peptide library was designed from GenBank phage ORFs (accessed June 1, 2018) using inclusion keywords (‘structural’, ‘tail’, ‘capsid’, ‘baseplate’, ‘holin’, ‘lysin’, among others) and excluding ORFs with keywords ‘initiator’, ‘chaperone’, ‘maturation protease’, and ‘morphogenesis’. ORFs were tiled into 56-aa peptides starting every 28 aa (50% overlap), deduplicated, reverse-translated using E. coli-optimized codons, and synthesized (Agilent SurePrint), yielding 277,051 oligopeptides. The library was cloned into a T7 phage display system (T7Select 415-1, Merck Millipore) using Liga5 (A&A Biotechnology) in place of T4 ligase with EcoRI and HindIII sites. Immunoprecipitation and sequencing were performed following a modified protocol based on Xu et al., 2015 (details in Supplementary Methods).

Sequencing reads were annotated using DIAMOND with BLASTx against a database built from the oligonucleotide FASTA. IgG-reactivity was quantified as raw reads per peptide; a protein was considered IgG-reactive if any peptide mapping to that protein had non-zero reads, and higher read counts indicated higher reactivity. Scheme presented in Figure 2, detailed description available in Supplementary Methods section. Library coverage relative to the metagenomic dataset was summarized post hoc by overlap between PhageScan-positive calls and metagenome-detected protein groups and host annotations (Supplementary Table S6).

**Figure 2.**
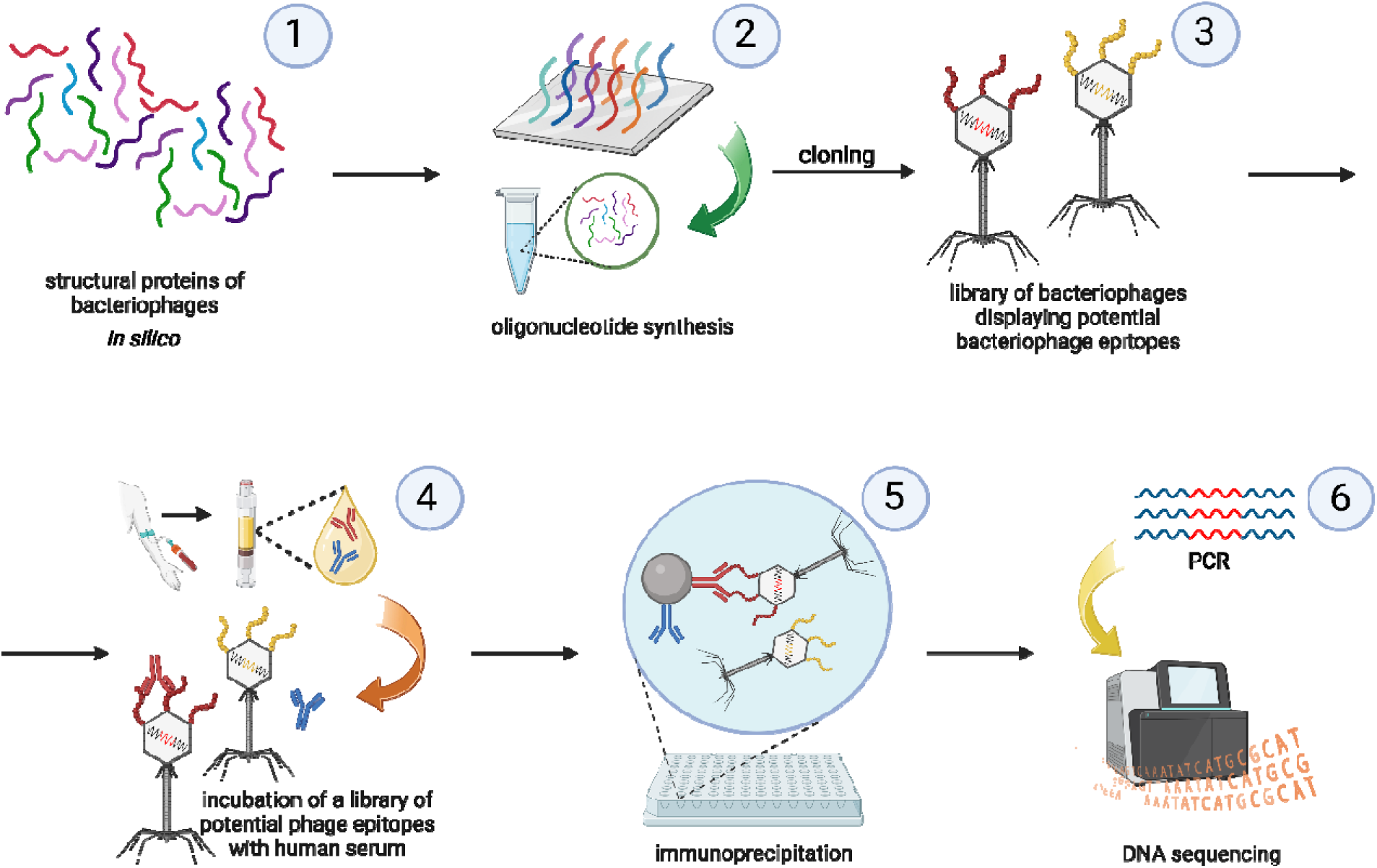
PhageScan technology (adapted from VirScan, Xu et al. (2015). Step 1: Structural protein sequences from bacteriophage genomes were computationally fragmented into overlap-ping 56-amino acid oligopeptides (28 aa offset). Step 2: Oligonucleotide sequences encoding 277,051 unique peptides were synthesized using SurePrint technology (Agilent). Step 3: Oligo-nucleotides were cloned into T7Select 415-1 vectors to generate a phage display library present-ing potential bacteriophage epitopes. Step 4: The amplified phage library was incubated with human serum samples to allow antibody-epitope binding. Step 5: Antibody-bound phages were captured using Protein A/G magnetic beads and isolated by immunoprecipitation. Step 6: En-riched phage inserts were PCR-amplified and identified by Illumina sequencing with subsequent DIAMOND annotation against the reference oligopeptide database. (created in https://BioRender.com)

### Translocation score

Translocation score was defined as an **intestine-to-serum ratio score**, computed as **log2((intestine+1)/(serum+1))**. Under this definition, **lower values indicate comparatively higher relative serum abundance**, whereas higher values indicate higher relative intestinal abundance. Associations between IgG reactivity (reads mapped to oligonucleotide database from phage-derived peptide library) and the translocation score were evaluated using Spearman correlation and a linear mixed-effects model with translocation score as the outcome and IgG reads as a fixed effect, including a random intercept for subject; the implementation additionally included a category variance component (0 + C(category)) when enabled in the analysis script. Model fitting used restricted maximum likelihood (REML) with the L-BFGS optimizer.

### Statistical analysis

All tests were two-sided unless otherwise stated. For comparisons of intestinal abundance between vOTUs classified as Translocated versus Only gut presence within taxonomic or host categories, we used Wilcoxon rank-sum tests on log10(TPM+1) values and controlled multiple testing using Benjamini-Hochberg false discovery rate (BH-FDR) within each analysis family (ICTV family set; host-class set). For protein-group analyses comparing Translocated versus Only gut presence within subjects, we computed per-subject medians and applied paired Wilcoxon signed-rank tests, with BH-FDR correction across protein groups. Effect sizes are reported as median differences (Δmedian) and the log2 ratio of medians (log2[median_trans/median_not]) alongside P values and FDR (Supplementary Table S3) With 37 paired subjects, the study has sensitivity primarily for moderate-to-large, paired effects; smaller effects may remain undetected, particularly after multiple-testing correction.

### Animal model of phage translocation

C57BL/6J male mice (n=8) were maintained under SPF conditions. Experiments followed EU directive 2010/63/EU, were approved by the 1st Local Committee for Experiments with the Use of Laboratory Animals (Wrocław, Poland; project no. 76/2011) and followed ARRIVE (Animal Research: Reporting of in vivo Experiments guidelines). T4 phage (ATCC) was propagated on E. coli B, purified by gel filtration followed by dialysis, and administered in drinking water (5×10 pfu ml ¹). After 26 h, gut content, intestinal mucus, mesenteric lymph nodes, and blood were collected and phage titers quantified by double-layer plaque assays. PBS-treated controls (n=3) were processed in parallel.

### Data and code availability

Sequencing Data is available in Sequencing Read Archive - PRJNA874475. Code for figures, and statistical analysis in available on github (https://github.com/olekszy/phage_transfer) IgG-reactivity library available upon request.

## Results

### In vivo phage retention across compartments

Translocation of particles from the intestine to the bloodstream is not a direct process but requires passage across the complex intestinal barrier. Particles must traverse mucosal layer, epithelial cells and underlying tissues, and are frequently transported via lymphatic pathways before entering the systemic circulation. We used a model phage T4 (Tequatrovirus, Straboviridae) to evaluate the extent to which phages can be retained during this passage *in vivo*. T4 phage (5×10 PFU/mL) (Plaque Forming Units/milliliter) was administered continuously in drinking water to mice (n=8) for 26 hours, and phage titers were quantified across four compartments: gut content, mucosal surface (those potentially contacting the transcitosing epithelium), phages in mesenteric lymph nodes (capturing material translocated by gut epithelial cells), and phages circulating in the blood, representing efficiently translocated ones.

Mean titers decreased from gut content (2.4×10 PFU/g) to intestinal mucus (2.4×10 PFU/g), mesenteric lymph nodes (8.7×10² PFU/g), and blood (2.7×10² PFU/mL) (Figure 3). This corresponded to stepwise reductions of ∼10-fold (gut content to mucus), ∼10-fold (gut content to lymph nodes), and ∼10-fold (gut content to blood). Blood titers were highly variable (median 20 PFU/mL, range 0-1250 PFU/mL), while lymph node titers remained over10-fold lower than gut content (Figure 3, Supplementary Figure S2). These findings indicate that the barriers encountered during translocation from the intestine lumen to the bloodstream exert a major constraint on phages, severely limiting the efficient translocation of functional viral particles.

**Figure 3.**
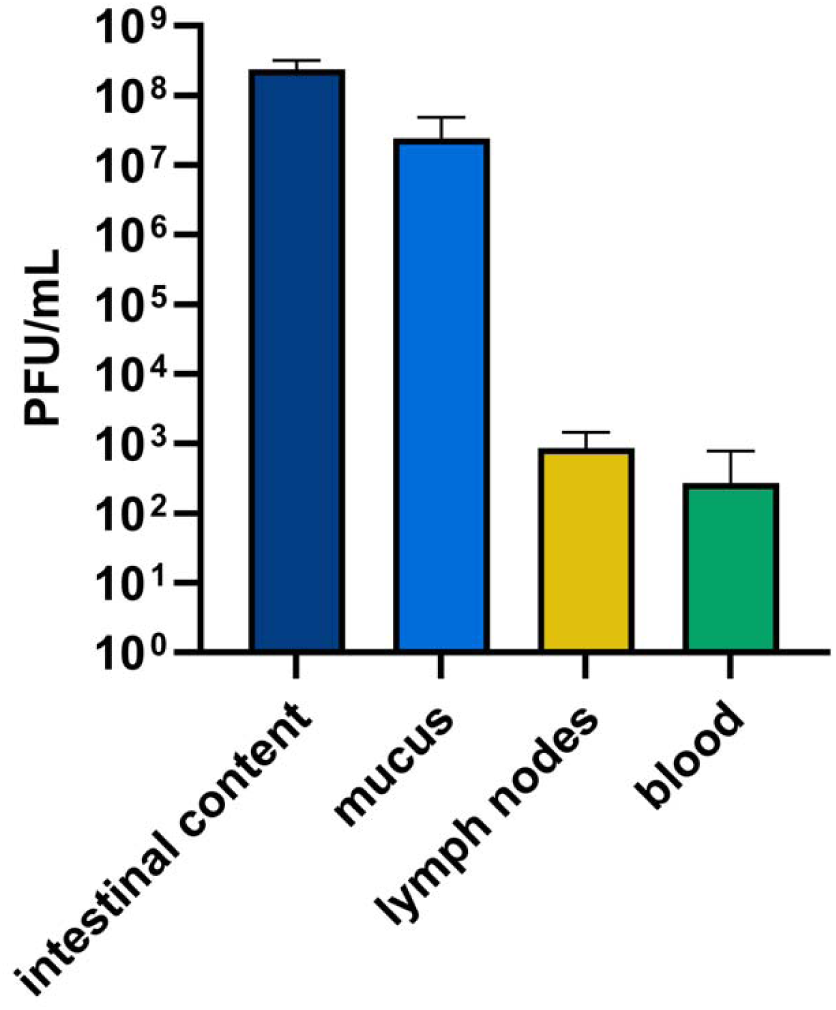
Mouse model of phage translocation and dissemination from gut. C57BL/6J normal male mice (N=8) were treated with Sepharose-purified T4 phage (ATCC) added to drinking water (5×109 PFU/mL) continuously for 26 hours. Murine blood, mesenteric lymph nodes, gut content, and scraped intestinal mucus were collected at 26 hours post-starting phage administration. Phage concentration was assessed by double-layer plate method or routine test dilution series. One exemplary experiment out of two conducted ones is presented.

Given the strong restrictiveness in translocation, we set out to identify which phages in natural phageomes are capable of crossing the intestinal-blood barrier, by characterizing the phageome present simultaneously in the gut mucosa and in systemic circulation.

### Intestinal mucosal viromes are dominated by Microviridae and filamentous phages

Phages present in the gut mucosa and in blood serum were identified in a group of 37 human patients. These individuals provided both colon mucosa biopsy samples and blood serum samples (Figure 1). Mucosal biopsies were used instead of gut content to exclude transient phages originating from food or water, which are unlikely to access the transcytosing endothelium. To focus on intact phage particles, virus-like particles (VLPs) were isolated, and only DNA encapsulated within VLPs was analysed, excluding DNA from disrupted capsids or lysed cells. This approach removed partially digested or degraded phage remnants that would not represent particles capable of translocation. Accordingly, each mucosal sample underwent CsCl gradient separation to isolate VLPs, followed by shotgun NGS sequencing.

Intestinal viral operational taxonomic unit (vOTU) abundance, normalized as transcripts per million (TPM), was highly skewed (0.66-7.76×10^7 TPM; median 40.2 TPM; 90% range 3.45-3.63×10^3; Figure 4A-D). Within phage family-assigned intestinal TPM, Microviridae (48.9%) and Inoviridae (42.4%) dominated, followed by Straboviridae (4.3%) and Herelleviridae (3.3%), with all remaining families contributing <1%. At the order level, assigned intestinal TPM mapped mainly to Petitvirales (53.2%) and Tubulavirales (46.1%), with Crassvirales contributing 0.75% (Figure 4C).

**Figure 4.**
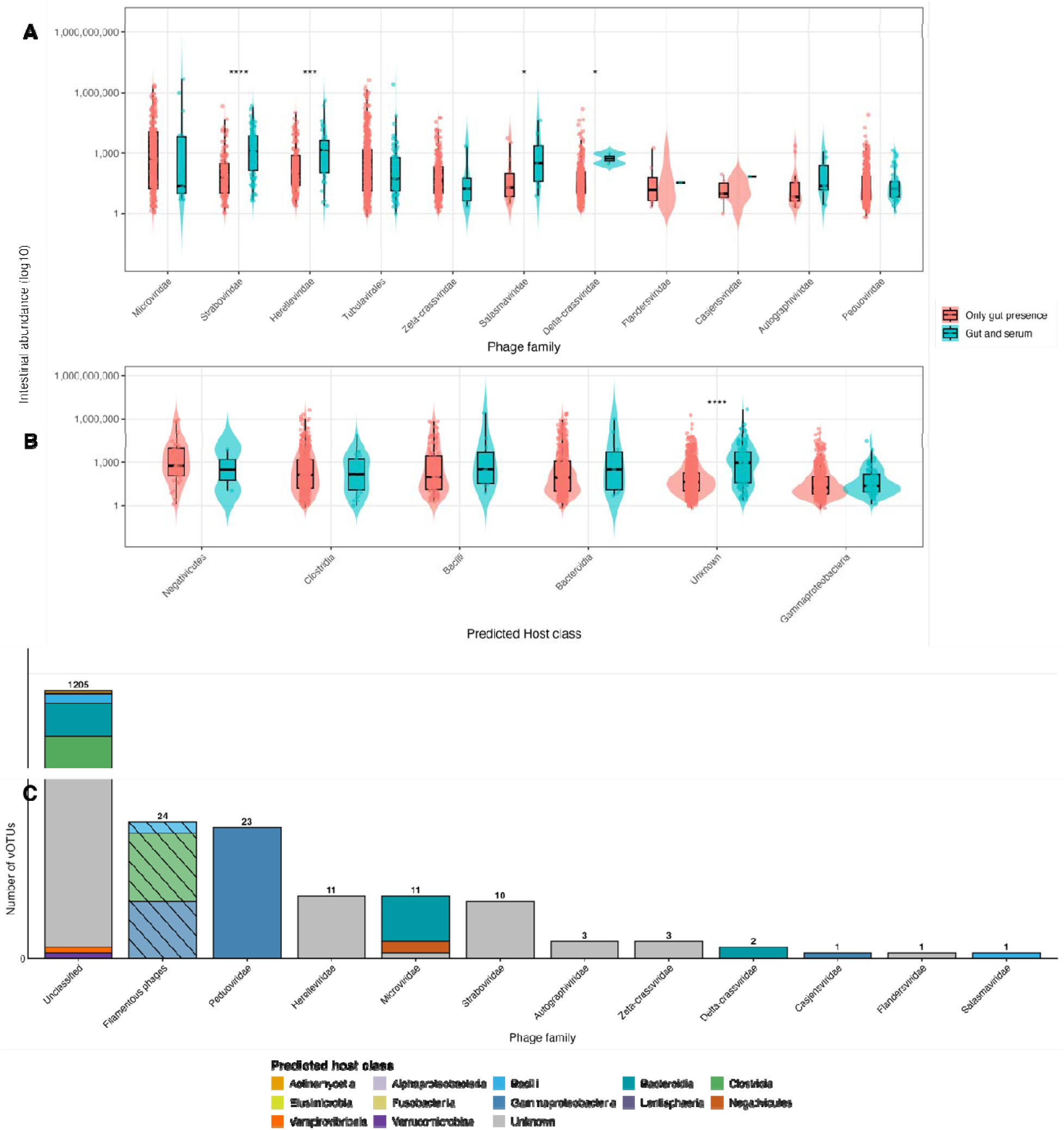
Taxonomic and host context of vOTUs detected in both intestine and serum. Intestinal abundance (log10[TPM+1]) for vOTUs grouped by ICTV family (A) or predicted host class (B), stratified by detection status within subjects: Translocated (abundance >0 in both intestine and serum) versus Only gut presence (intestine >0, serum =0). Points represent individual vOTUs; boxplots show the median (center line), interquartile range (box), and 1.5×IQR whiskers. Two-sided Wilcoxon rank-sum tests compare Translocated versus Only gut presence within each category, with Benjamini-Hochberg false discovery rate (BH-FDR) correction across categories; n values per group are shown on the plot and summarized in Supplementary Table S3. (C) Stacked bar plot of translocated only vOTU counts per family (axis break), filled by predicted host class; totals are labeled. vOTU, viral operational taxonomic unit.

### Microviridae and Tubulavirales dominate translocated phage abundance

Translocation within matched gut-serum samples was defined as the detection of the same vOTU in both compartments. Interestingly, translocation was not universally associated with higher intestinal abundance across all phage vOTUs - after BH-FDR adjustment, elevated gut abundance among translocated observations was limited to a specific subset of families. In absolute terms, Microviridae accounted for the largest share of translocated phage abundance, consistent with their overall dominance in the mucosal virome. However, their intestinal abundance did not differ significantly between translocated and gut-only vOTUs (FDR=0.5, Table 1). This indicates that translocation for Microviridae occurs irrespective of their local abundance in the mucosa. On the other hand, after BH-FDR correction, Straboviridae and Herelleviridae showed significantly higher intestinal abundance among translocated versus gut-only observations (Table 1; Supplementary Table S3), suggesting that for these families translocation may be based on the higher local abundance at the mucosal surface.

**Table 1.** summary table of the per-phage family Wilcoxon rank-sum tests comparing intestinal abundance (“presence”) between two groups of vOTUs: those classified as Translocated (detect-ed in both gut and serum within a subject) versus Only gut presence (detected in gut but not se-rum). *Filamentous phages presented as Tubulavirales.

| Phage Family | Median presence in translocated (TPM) | Median presence in gut only (TPM) | pvalue | FDR correction |
| --- | --- | --- | --- | --- |
| Microviridae | 23.03 | 477.61 | 0.32 | 0.50 |
| **Straboviridae** | **1295.77** | **60.17** | **1.41E-10** | **1.55E-09** |
| **Herelleviridae** | **1434.70** | **87.25** | **9.88E-04** | **0.0054** |
| *Tubulavirales\** | 52.25 | 98.77 | 0.22 | 0.44 |
| Zeta-crassviridae | 17.37 | 43.51 | 0.37 | 0.51 |
| Salasmaviridae | 317.30 | 19.56 | 0.05 | 0.13 |
| Delta-crassviridae | 621.36 | 31.25 | 0.05 | 0.13 |
| Flandersviridae | 34.81 | 19.32 | 0.89 | 0.89 |
| Casjensviridae | 68.97 | 9.52 | 0.50 | 0.60 |
| Autographiviridae | 23.10 | 6.74 | 0.24 | 0.44 |
| Peduviridae | 17.40 | 15.18 | 0.54 | 0.60 |

Among phage family-assigned translocated abundance, Microviridae and Tubulavirales (filamentous phages were analyzed at the order level) dominated (8.69×10^6^ TPM combined; Figure 4A,C,D). For Straboviridae and Herelleviridae, translocated contigs were associated with significantly higher intestinal abundance compared with contigs observed only in the intestine. (medians 1,296-1,435 vs 60-87 TPM; FDR<0.01; Figure 4A, Table 1). Host-associated patterns revealed 21-fold higher abundance of unknown-host vOTUs in translocated ones than in those observed only in gut (FDR=2.03×10^−17; Figure 4B).

To avoid distortion of detected viral composition by phi29 whole-genome amplification (WGA), we performed a computational assessment of length-dependent bias. Across subjects, vOTU contig length was not inversely associated with intestinal abundance; instead, within-subject Spearman correlations between log10(TPM+1) and log10(length) were weakly positive (median ρ = 0.156; IQR 0.117-0.215; 33/37 subjects positive; Supplementary Table S4). In addition, Microviridae vOTUs were not enriched among the shortest sequences (median length 5,786 bp versus 3,436 bp for other annotated families; only 10.9% of Microviridae fell in the shortest-length quartile; Supplementary Table S4). Together, these patterns are not consistent with a dominant global length-driven amplification artifact explaining Microviridae prominence.

### Functional associations of translocated protein coding regions

Protein-coding genes were predicted, and functional annotation was performed to identify potential function-related patterns associated with mucosal and translocating phages. Among module-assigned structural proteins, 32% of CDS records (923/2,861) were detected in both intestine and serum (Figure 5A). Translocation frequency, defined as fraction of contigs co-detected in paired samples, ranged from 7.8% (lysis) to 60.2% (baseplate), with intermediate frequencies for tail (24.9%), capsid (40.1%), and other structural groups (36.3%; Figure 5A). No function reached false discovery rate (FDR) significance (minimum q=0.521; Figure 5A); overall, the most frequently observed functions of translocated phage genomic content were dominated by structural annotations (e.g., structural protein, tail fiber protein, capsid/scaffold, and baseplate) (Supplementary Table 1).

**Figure 5.**
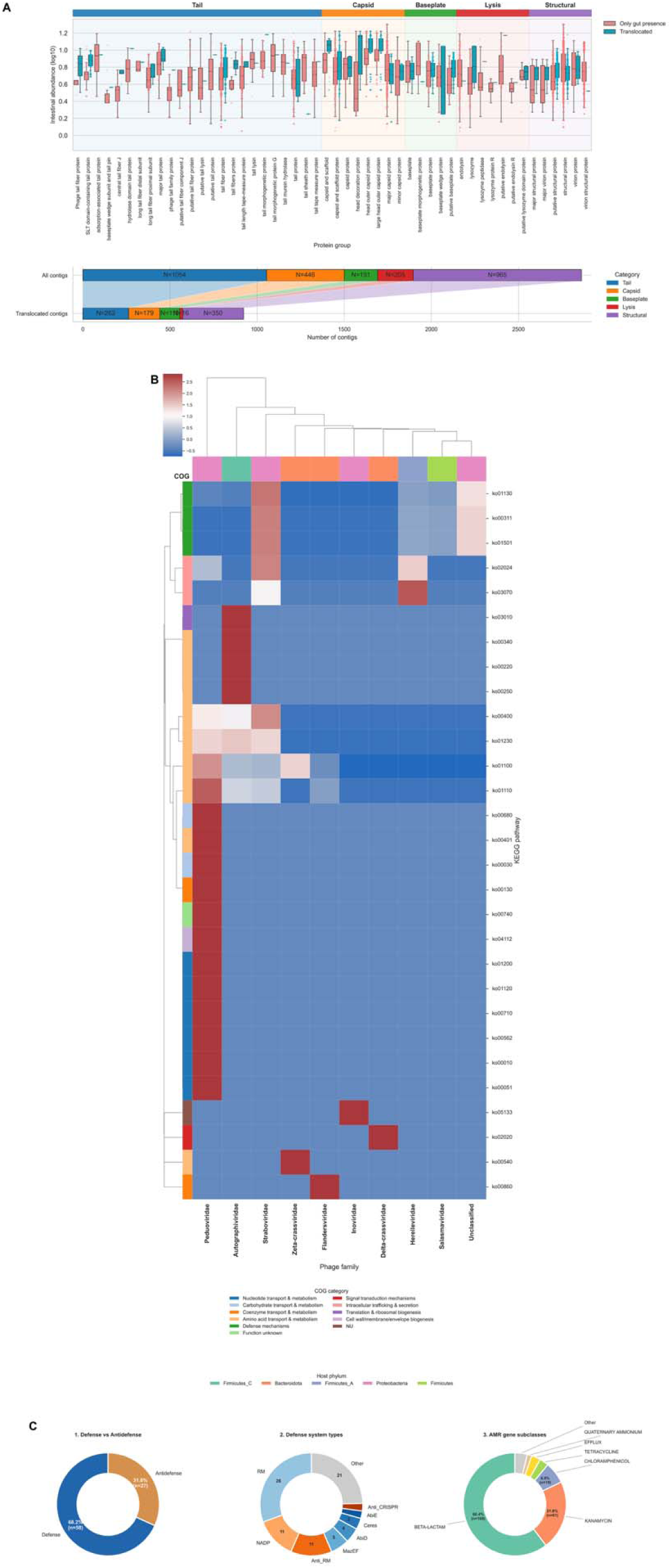
Functional and defense signatures among translocated phage contigs. (A) Intestinal abundance distributions (log10-transformed) of protein groups detected in the intestine, stratified by whether the corresponding contigs were also detected in serum (Translocated) or were restricted to gut detection (Only gut presence). Protein groups are ordered and visually grouped by predicted structural module (Tail, Capsid, Baseplate, Lysis, Structural; colored header bar), with shaded backgrounds indicating module blocks. Points represent per-sample values and boxplots summarize the distribution across samples; significance markers (asterisks) indicate multiple-testing-corrected paired differences between Translocated and Only gut presence within each protein group. The lower subpanel summarizes contig counts per structural module for all contigs versus translocated contigs, highlighting shifts in module representation between the full set and the translocated subset. (B) Heatmap of KEGG pathway enrichment patterns across Phage families, computed as row-wise Z-scores of total abundance aggregated per KEGG pathway × phage family. Rows (KEGG pathways) are annotated by the dominant COG functional category and columns (phage families) by the dominant predicted host phylum. Hierarchical clustering (dendrograms) organizes pathways and families by similarity in their Z-score profiles. Legends report the COG-category and host-phylum color mappings. (C) Summary of accessory functional potential detected among translocated contigs, shown as three donut charts.1), the proportion of contigs carrying D**e**fense versus Antidefense signatures (percentages with counts). 2) the distribution of the top defense system types across contigs. 3) the distribution of AMR gene subclasses across contigs

Across 37 matched intestine-serum pairs, 29,733 protein-coding genes showed gut-only detection (intestine present, serum absent), spanning 301 protein groups; these counts reflected CDS calls pooled across subjects (Figure 5). Most gut-only sequences were annotated as unknown function (86.1%), leaving 4,126 annotated proteins (14%). Among annotated gut-only protein coding genes, DNA/RNA/nucleotide metabolism (33.5%) and moron/auxiliary metabolic gene and host takeover (18.5%) were most common, followed by head/packaging (7.9%) and integration/excision (7.0%). Gut-only abundance summaries (log10[TPM]) highlighted frequently detected replication- and toxin-associated functions, including replication initiation protein (36/37 subjects; median 0.952) and a RelE-like toxin (35/37; 0.850), alongside structural categories such as major head protein (36/37; 0.628) (Supplementary table 1).

Translocated contigs with EggNOG-derived annotations mapped to 29 Kyoto Encyclopedia of Genes and Genomes (KEGG) orthology (KO) pathway identifiers across 10 phage families (102 pathway-family entries) in the KO-assigned subset (Figure 5B). Across this subset, the highest-abundance pathway identifiers were led by biosynthesis of antibiotics (ko01130), beta-lactam resistance (ko01501), and penicillin and cephalosporin biosynthesis (ko00311), followed by metabolic pathways (ko01100) and biosynthesis of secondary metabolites (ko01110) (Figure 5B). Straboviridae contributed the largest share of KO-assigned abundance (2,053.6 TPM; 36% of the KO-assigned total), and Clusters of Orthologous Groups (COG) categories were dominated by COG E (amino acid transport and metabolism) and COG V (defense mechanisms), together accounting for 58% of pathway-family entries (Figure 5B). DefenseFinder analysis identified 85 defense-related systems (comprising 142 gene hits) across the vOTU representatives, of which 58 were classified as defense and 27 as anti-defense. Restriction-modification (RM; n = 26) and NADP-associated systems (n = 11) were the most prevalent types. Putative AMR subclass annotations were detected in 280 contigs, dominated by beta-lactam-associated genes (n = 169; 60.4%) and aminoglycoside-associated genes (kanamycin subclass; n = 61; 21.8%) (Figure 5C, Supplementary Table 2). This is consistent with a growing body of evidence indicating that bacteriophages can acquire and carry AMR gene (Liao et al. 2024). Thus, beyond virion structural proteins, translocated phage contigs retained measurable accessory annotations among AMR subclasses and defense/anti-defense gene homologs, highlighting that serum-detected phage sequences included resistance-associated features (Niault et al. 2025) suggesting that gut-to-blood phage translocation may constitute a route for disseminating resistance and phage-defense determinants to bacterial communities beyond the intestinal compartment.

### IgG-recognition of phages versus compartment detection and translocation score

Due to significant homology between some phage proteins, antibodies induced by one phage often cross-react with another phage (Gembara & Dąbrowska, 2021). To avoid ambiguous results, we analyzed immune responses to translocating phages at the protein level. We applied a modified VirScan approach (Xu et al., 2015) with a phage-derived epitope library containing 277,051 oligopeptides that cover the structural proteomes of phages available in GenBank (see Materials and Methods and Figure 2 for details). Briefly, the library was immunoprecipitated with each patient’s serum, and massively sequenced to identify phage structural oligopeptides recognized by the patients’ antibodies.

In a population-level analysis, where all samples were analyzed collectively, immunoglobulin G (IgG) reactivity showed weak associations with the intestine-to-serum ratio score (log2((intestine+1)/(serum+1))). Lower numbers of reads in IgG-reactive fraction of epitopes were associated to higher relative serum abundance of phages bearing those epitopes (phage host: ρ=-0.26, p=0.093; lifestyle: ρ=-0.09, p=0.57; protein function: ρ=0.14, p=0.20; Figure 6D). Some IgG-reactive protein groups, to some extent, overlapped with compartment-detected groups (32/163 intestine-detected; 26/95 serum-detected), but most groups recognized by IgGs in the investigated sera (102/134; 76.1%) lacked metagenomic detection in either compartment (Figure 6A) suggesting, that majority of phages that can be targeted by specific human IgGs are not present in the gut mucosa nor serum.

**Figure 6.**
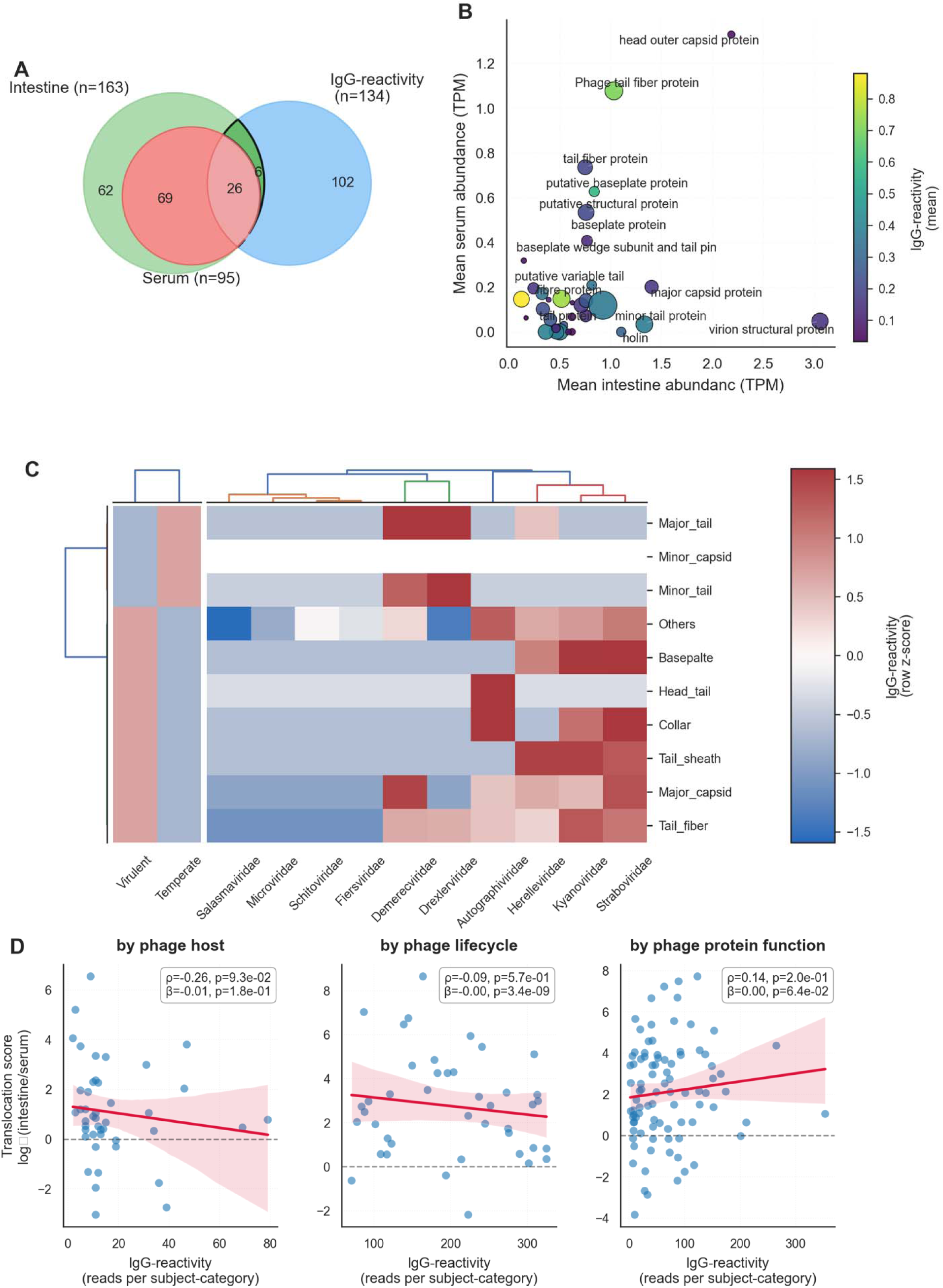
IgG-reactivity across gut-serum phage proteins and association with translocation. (A) Venn diagram showing overlap of protein groups detected in intestine, serum, and IgG-reactivity datasets (presence defined as non-zero signal) (B) Protein-group mean intestine abundance (x-axis) versus mean serum abundance (y-axis); point colour indicates mean IgG-reactivity (raw counts) and point size indicates total IgG-reactivity (log10-scaled sum), with labels shown for the highest IgG-reactive groups. (C) Dual clustered heatmaps of transformed IgG-reactivity reads (log1p, row z-score) summarised by phage cycle (left) and phage family (right), sharing a row dendrogram. (D) Intestine-to-serum ratio score log2((intestine+1)/(serum+1)) versus IgG-reactivity (raw reads per subject-category), stratified by phage hosts, predicted lifecycle, and protein function; lower scores indicate comparatively higher relative serum abundance. Inset statistics report Spearman’s ρ and mixed-effects model slope (β).

Among the 26 functional groups detected in the intestine and serum and targeted by specific IgGs, tail fiber proteins showed detectable IgG reactivity and were frequently detected in serum or intestinal mucosa (Figure 6B). Those with detectable IgG reactivity but lower presence in serum and intestinal mucosa fell into categories of putative variable tail fiber protein and tail protein. IgG-reactive phage epitopes that did not correspond to any phage material detected in serum or intestine (Figure 6A) included strongly recognized tail-fiber categories (e.g., putative short tail fibre, tail fiber like protein, putative variable tail fibre protein, tail protein) (Figure 6B). Taking primary function of tail-fibers, this may reflect a direct neutralizing effect of antibodies targeting phage proteins required for recognition of and propagation on the relevant bacterial host.

When stratified by predicted lifestyle, minor tail/capsid and major tail functions showed higher aggregated IgG-reactivity in temperate predictions, whereas collar, tail sheath, major capsid, head-tail and baseplate-associated labels were higher in virulent predictions (Figure 6C). To quantify how well our peptide library recapitulated signals observed in the current metagenomic vOTU set, we summarized overlap (“sensitivity”) at multiple annotation levels. Among metagenomically detected protein groups, 32/163 (19.6%) were also detected by PhageScan; at the predicted host genus level, 4/132 (3.0%) overlapped; and at normalized host labels, 14/109 (12.8%) overlapped (Supplementary Table S6).

Notably, this population-level analysis may be influenced by the high heterogeneity of individual serological profiles and phageome composition, therefore we extended this analysis with individual mapping of epitope recognition in patients.

### Phages recognized by specific IgGs were absent from gut and serum phageomes in 90% of individuals

To quantify the relationship between humoral anti-phage responses and the composition of the phageome, we mapped the full epitope library (277,014 oligopeptides) to each patients’ metagenomically detected phageome in gut mucosa and serum and then compared it with the individual IgG-reactive (immunoprecipitated) subset of epitopes. Per patient, 6-449 library oligopeptides (median 141; 0.002-0.162% of the library) matched phageome-detected sequences, representing the maximum set of epitopes each individual could recognize given their current phageome. Immunoprecipitation identified 90-245 IgG-reactive epitopes per patient (median 147), yet these overwhelmingly corresponded to phage proteins absent from the patient’s own virome. Across 30 individual, only 3 patients showed any overlap between the immunoprecipitated and virome-matched epitope sets; while the remaining 27 patients showed zero overlap. When expressed as a fraction of IgG-targeted epitopes, 0-2.08% of immunoprecipitated hits corresponded to epitopes present in the patient’s virome (median 0%). Conversely, when expressed as a fraction of virome-matched epitopes, patients mounted detectable IgG responses against 0-0.78% of the phage epitopes present in their virome (median 0%).These results indicate that circulating anti-phage IgG is directed almost exclusively against epitopes not represented in the individuals’ current intestinal of circulating phageome, that is, no longer detectable in the sampled virome.

### Sequencing dark matter among translocated vOTUs and sequence-level signatures

Translocated vOTUs were dominated by sequencing dark matter: 1,205/1,295 (93.1%) lacked a confident assignment leaving 90 vOTUs as taxonomically “known” anchors (Figure 7A, D). Unassigned vOTUs nonetheless formed reproducible co-enrichment links to phage-assigned vOTUs in the serum association network, with connectivity concentrated around Herelleviridae, Straboviridae, Peduoviridae, and Inoviridae (each connected to 90-124 unique dark-matter partners; Figure 7A, D). Hub-centered wiring indicates that serum translocation involves repeatable phage modules in which a few annotated families act as anchors for a much larger, unclassified virome.

**Figure 7.**
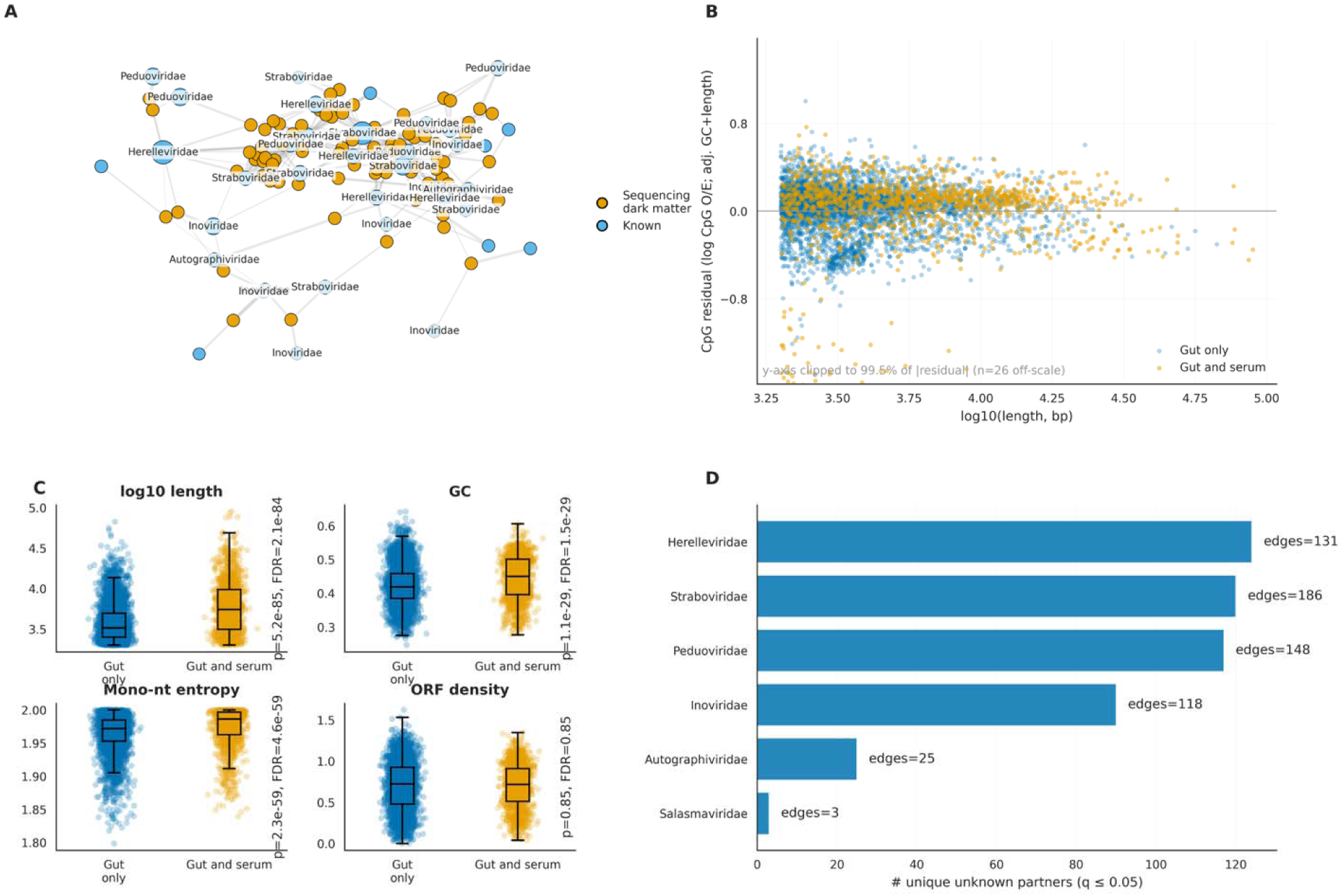
Co-translocation association network and sequence signatures of translocated viral contigs. (A) Unknown-known co-enrichment network of serum-enriched viral contigs across subjects (n=37). Nodes represent vOTUs and are colored by annotation class: Sequencing dark matter (contigs lacking a confident assignment) and Known (confidence ≥70). Serum enrichment is defined per subject as serum abundance ≥1.0 and log2((serum+0.5)/(intestine+0.5)) ≥1.0. Edges connect unknown-known pairs that co-occur above expectation; significance is assessed by one-sided hypergeometric tests and corrected by BH-FDR (q ≤ 0.05), with additional thresholds for co-occurrence (≥3) and association strength (phi ≥ 0.25). Node size scales with degree and edge width scales with phi. (B) CpG residual versus vOTU length (log10 bp), colored by annotation class; outliers are clipped for display as indicated. (C) vOTU length comparison for Translocated versus Only gut presence groups. (D) Comparisons of GC content, mono-nucleotide Shannon entropy, and ORF density between Translocated and Only gut presence vOTUs. For panels C-D, two-sided Mann-Whitney U tests are used with BH-FDR correction across tested features; P and FDR values are shown on the plots. Boxplots show median, interquartile range, and 1.5×IQR whiskers.

CpG residuals (observed/expected ratio after GC and length adjustment) varied widely, with 26 outliers clipped for display (Figure 7B). Sequence properties differed between translocated and intestine-only vOTUs: translocated vOTUs were longer (median 5,508 vs 3,265 bp; p=5.2×10), had higher GC content (45.0% vs 41.9%; p=1.1×10 ²), and higher mono-nucleotide Shannon entropy (p=2.3×10), while open reading frame (ORF) density did not differ (p=0.85) (Figure 7C). Protein annotation remained incomplete, with 63.2% of translocated protein-coding sequences labeled “hypothetical,” and the most frequent specific labels mapping to virion structural components (e.g., tail fiber, baseplate, head proteins).

## Discussion

### Filtering gut-to blood phage translocation

Profiling of gut phage translocation across matched intestinal-serum samples is consistent with animal model of T4 phage (Figure 4) and indicates that systemic detection reflects strong anatomical filtering rather than simple spillover from intestine. Across the cohort, phage abundance decreased by ∼98% from intestinal mucosa to serum (Figure 4A-B), and translocated vOTUs were dominated by unannotated sequences (93.1%; Figure 7A, D) The *in vivo* tracer model revealed a similar pattern: orally administered T4 showed stepwise loss across compartments, with blood titers approaching the detection limit even after continuous exposure (Figure 3). These findings suggest that translocation is determined not only at the epithelial barrier but also by downstream processes such as lymphatic filtration and systemic clearance (Berkson et al. 2024; Dąbrowska 2019). Unexpectedly, and contrary to observations from *in vitro* systems (Nguyen et al. 2017), mucosal surfaces *in vivo* did not appear to efficiently accumulate phages. Instead, an approximately one-order-of-magnitude reduction in phage abundance was observed during passage from the gut lumen to the mucosal surface.

Direct quantitative comparison between phage detected in lymph nodes and phage reaching the circulation is difficult. Interpretation is complicated by several factors, including the unknown rate of phage neutralization within lymphatic tissue, dilution in blood serum, and deposition in other tissues and organs after entry into the circulation. However, these data together offer a parsimonious explanation for why serum viromes are reproducibly detectable yet remain orders of magnitude below gut reservoirs. Low level epithelial entry along with cumulative loss at successive barriers limits persisting phage detectability in blood.

### Taxonomic and functional determinants of translocation

A substantial fraction of the mucosal virome (42.4% of family-assigned abundance) mapped to filamentous phages within the order of Tubulavirales. We report filamentous phages detections at the order level rather than as Inoviridae, because current ICTV taxonomy restricts Inoviridae to phages of Gram negative bacteria (Knezevic, Adriaenssens, and Ictv Report Consortium 2021). Metagenomic databases assign the “Inoviridae” label based on computational detection tools (Roux et al. 2019) that encompass phylogenetically distinct lineages infecting Firmicutes, which share no sequence identity with any classified Inoviridae member (Burckhardt et al. 2023) and likely represent novel, as-yet-unclassified families within Tubulavirales (Billaud, Petit, and Lossouarn 2023).

At the intestinal mucosa, the virome was dominated by Microviridae (48.9%) and Tubulavirales (42.4%) (Figure 4A-D), families that are well represented in gut phage catalogues and are frequently detected across cohorts. Filamentous phages proliferate without lysing host cells and can affect host fitness, aligning with the pronounced partitioning of filamentous phages between Firmicutes classes reported herein (Figure 4B-D). By contrast, Crassvirales contributed 0.75% (Figure 4C), despite their reported abundance in many gut studies, suggesting that sampling context (for example, mucosal versus luminal) and/or cohort features may shift the balance among dominant phage groups (Smith et al. 2023; Zuo et al. 2019).

Functional annotations differed between coding sequences detected only in intestinal virome and the subset showing compartment overlap. Most coding sequences were detected only in intestinal virome (29,733 proteins; 301 groups; Figure 5). A smaller subset showed matched intestine-serum co-detection, with structural module CDS represented among overlapping detections (32% of structural CDS detected in both compartments; Fig. 4A). Because all phages encode core structural genes, this pattern likely reflects which structural CDS were detectably recovered in each compartment, rather than capsid genes being missing from gut-only phages. It is consistent with experimental evidence that phage virions can cross epithelial layers by transcytosis (Nguyen et al. 2017). Variation across structural modules (baseplate 60.2%, capsid 40.1%, tail 24.9%, lysis 7.8%; Figure 5A) is compatible with differences in detectability and annotation across module-associated CDS between compartments, consistent with reports that downstream virome functional screens apply category-dependent curation for structural and attachment-related genes (Ramírez et al. 2026). However, direct mechanistic interpretation will require experiments that test whether virion architecture and surface exposure influence uptake or trafficking.

Functional enrichment signals among translocated contigs (Figure 5B-C) should be interpreted conservatively. In particular, KEGG terms linked to antibiotic resistance and the putative presence of AMR-like subclass hits (Figure 5C) do not, on their own, establish phage-encoded resistance, given evidence that virome pipelines frequently overcall ARGs and that many apparent hits reflect false positives or non-resistance homologs (Enault et al. 2017). With those factors in mind, the accessory annotations detected on translocated contigs – putative AMR determinants, restriction modification systems, and anti-defense gene homologs –gain confidence from the multistep VLP purification that preceded annotation: CsCl ultracentrifugation, 0,22 µm filtration, DNase treatment, host-read depletion collectively minimize the bacterial DNA carryover that drives false-positive ARG calls in virome studies (Enault et al. 2017; Billaud et al. 2021). Prophage-encoded ARGs can be functionally mobilized to confer resistance in heterologous hosts (Liao et al. 2024; Pfeifer, Bonnin, and Rocha 2022), and phage-borne defense systems are actively disseminated by lateral transduction (Rousset et al. 2022; Kuang et al. 2026). Because these elements were detected on contigs present in serum, gut-to-blood phage translocation may constitute a route for delivering resistance and defense determinants to extra-intestinal bacterial communities-a hypothesis that should be tested by direct validation in mammalian models.

### Dark matter dominance and sequence signatures of translocated vOTUs

An important part of the paired-compartment analysis is that most translocated vOTUs lacked confident database assignment (93.1%; Figure 7A,D), mirroring the wider “viral dark matter” problem in gut viromics (Fitzgerald et al. 2021; Santiago-Rodriguez and Hollister 2022). Network structure provided reproducible anchors: taxonomically assigned families formed connectivity hubs linked to many dark-matter partners (Figure 7A,D), suggesting that co-enrichment relationships can give background for unassigned sequences when taxonomic resolution is limited among functionally related groups even when precise taxonomy is absent (Fitzgerald et al. 2021; Santiago-Rodriguez and Hollister 2022). Sequence features further separated vOTUs detected in gut and serum from those that were gut only (Figure 7B-C), including shifts in contig length and GC content (Figure 7C) and a modest but consistent elevation in CpG residuals after GC- and length-adjustment (Wilcoxon rank-sum/Mann-Whitney U; median = 0.0429, BH-FDR q=3.28×10^-11^;Figure 7B). Because CpG content and context influence immunogenicity, nucleotide composition may correlate with systemic detectability of phage DNA. Bacteriophage genomes are derived from bacteria and therefore contain largely unmethylated CpG motifs. Such unmethylated CpGs are potent activators of the innate immune system via TLR9, stimulating dendritic cells, B cells, and pro-inflammatory cytokine production. It remains unclear how such stimulation might influence phage translocation. Activated (by phage CpG) epithelial cells may enhance endosomal trafficking and increase surface receptor expression, so we hypothesize that this activation could contribute to higher transcytosis. However, the balance between enhanced or disrupted cellular transport, as well as between transport and degradation, particularly by activated immune cells, is extremely complex, and this issue requires further investigation (Kharrat et al. 2025).

### Immune recognition and clearance dynamics

We evaluated the overall association between phage-derived epitope recognition by IgGs in the studied cohort and the phages detected in their gut or serum viromes. At the population level, this analysis revealed a weak negative association between IgG reactivity and the translocation score (Figure 6D). Also, most IgG-reactive protein groups lacked metagenomic detection in either compartment (Figure 6A).

Because IgG recognition is highly specific-different individuals may recognize particular phage epitopes without affecting the same phages in others-we also analyzed epitope recognition at the individual patient level. In 90% of individuals, phages recognized by specific IgGs were absent from their gut and serum phageomes. Among the remaining 10%, the largest fraction of phageome epitopes detectable in a single patient (considering library capacity and the actual phages present) that were recognized by IgGs reached only 0.78%. This is consistent with durable humoral memory (Kaźmierczak et al. 2021; Brzozowska et al. 2023; Gembara and Dąbrowska 2021) and compatible with partial neutralization and accelerated clearance of opsonized phage material preventing it from accumulation in serum (Berkson et al. 2024). These dynamics have practical relevance: cellular uptake of antibody-opsonized phages and phage-neutralizing immunity can act as sinks that reduce systemic phage persistence (Bichet et al. 2021) and may translate into differences in observed translocation efficacy between phages.

Importantly, the high abundance of phages in the gut mucosa suggests that most phages benefit from mucosal tolerance and are not targeted by mature, neutralizing antibodies. This aligns with general patterns observed in mucosal microbiota: even when microbes are targeted by secretory IgA, they are typically prevented from directly adhering to the gut epithelium and remain anchored within the mucus layer, rather than being efficiently destroyed (Chen et al. 2020; Rogier et al. 2014; MacPherson et al. 2018). In contrast, phages that have elicited mature, specific immune responses-manifested by detectable serum IgG levels, which can also be present in the mucosa- are rapidly cleared, disappearing not only from the circulation but also from the gut mucosa.

### Limitations and outlook

This study is constrained by factors that commonly limit mechanistic inference in metagenomic viromics. High fractions of unclassified vOTUs and hypothetical proteins (Figure 7; Figure 5) restrict the identification of molecular determinants of epithelial uptake and immune recognition.

In addition, short-read sequencing data cannot always distinguish integrated prophage DNA from encapsidated genomes, limiting direct inference of phage life cycles. In serum, differences in recovery of DNA versus whole virions, together with rapid immune clearance can further contribute to non-detection. Future studies should incorporate extraction blanks and synthetic spike-in standards processed alongside samples to quantify cross-sample carryover and establish sensitivity limits for low-biomass serum viromes. In the in vivo tracer model that focuses on a single phage and defined exposure window, the framework should be extended to broader phage diversity to validate generalization.

Mechanistic dissection using isolate panels and epithelial/colonoid systems can test whether particle architecture, surface properties, and receptor-binding modules predict translocation efficiency (Nguyen et al. 2017; Le et al. 2024).Focused functional validation-particularly of dark-matter proteins and ambiguous metabolic/AMR-like annotations-will be essential to connect sequence features to translocation competence and immune outcomes (Enault et al. 2017; Santiago-Rodriguez and Hollister 2022).

## Conclusions

This study demonstrates that phage signals identified in blood are primarily indicative of very limited passage through the intestinal barrier, followed by subsequent systemic elimination, rather than direct leakage from gut levels. By coupling matched colon mucosa-serum metagenomics with IgG profiling and an oral T4 tracer model, we outline a stepwise filtration framework in which only a small, compositionally biased fraction of gut phage material reaches-and persists in-circulation. As summarized in Figure 8., high diversity and amounts of phages found in mucosa suggest that phages widely benefit from mucosal tolerance (Step 1); we further propose that low-frequency epithelial passage (Step 2) is followed by lymphatic redirection to draining lymph nodes (Step 3), a checkpoint that may promote phage-specific response maturation and specific IgG production (Step 4). Phage-specific IgG dramatically reduces serum detectability of phages, with both systemic and gut persistence limited by accelerated neutralization and clearance (Step 5a and b).

**Figure 8.**
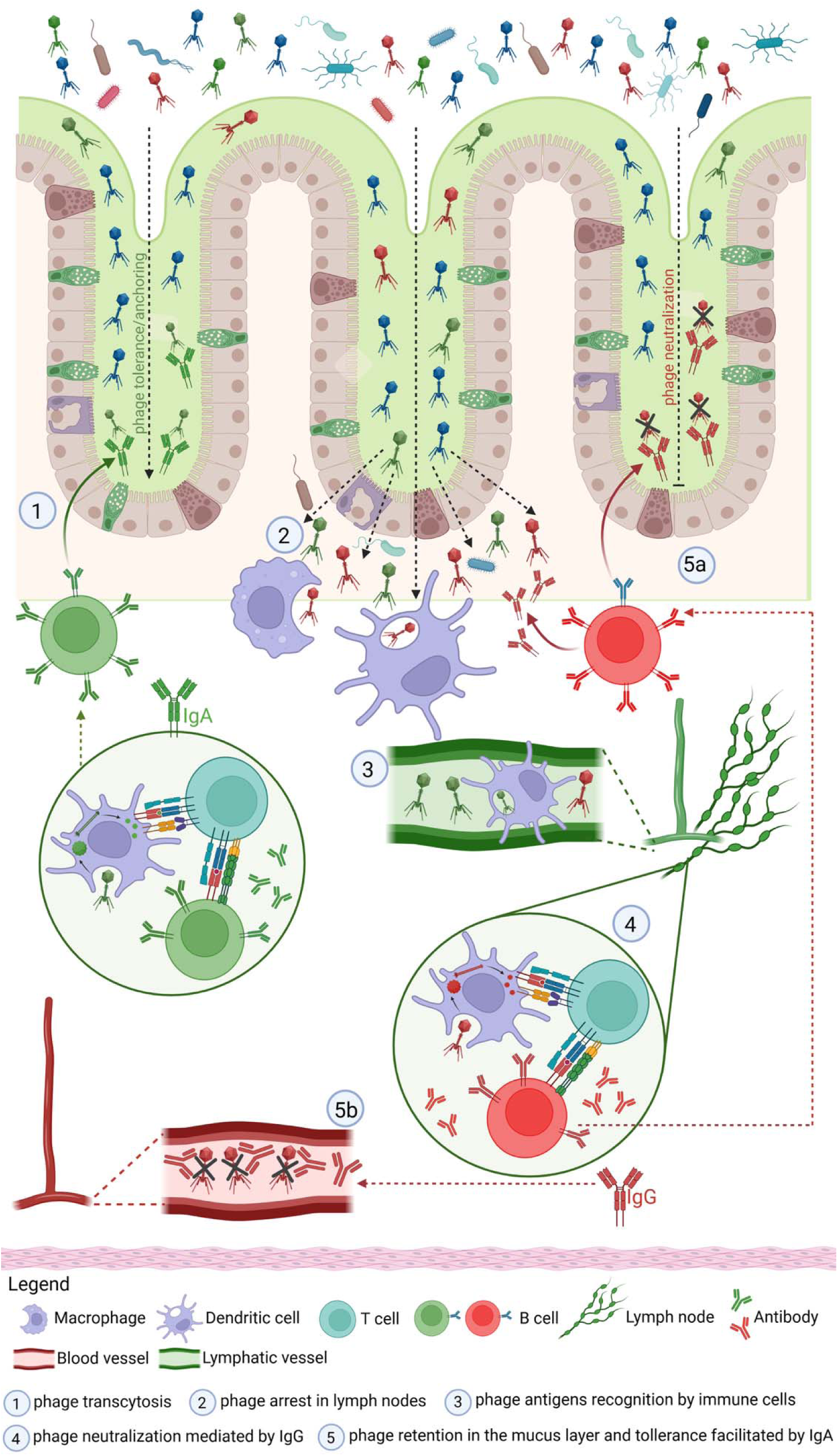
Proposed immunological mechanism regulating phage translocation. (1.) Phages commonly benefit from mucosal tolerance; (2.) their low-frequency epithelial passage (3.) is followed by lymphatic redirection to draining lymph nodes; (4.) phage-specific response maturation and specific IgG production result in (5.) dramatic reduction of phage survival both systemic and in gut mucosa (created in https://BioRender.com).

## Acknowledgments

This work was supported by National Science Center (Poland) grants: Preludium16 UMO-2018/31/N/NZ6/02584, Opus18 UMO-2019/35/B/NZ7/01824, and by the subsidy for scientific activity provided to the Department of Molecular Microbiology by the University of Lodz (B2111000000038.01).

## References

Almeida, Gabriel M. F., Elina Laanto, Roghaieh Ashrafi, and Lotta-Riina Sundberg. 2019. “Bacteriophage Adherence to Mucus Mediates Preventive Protection against Pathogenic Bacteria.” mBio 10 (6): e01984–19.

Badillo-Pazmay, Gretta Veronica, Carlo Fortunato, Laura Cianfruglia, Federica Novazzi, Pietro Giorgio Spezia, Luigi Rosa, Dolores Limongi, et al. 2025. “The Gut and Circulating Virome: Emerging Players in Aging and Longevity.” Frontiers in Aging 6: 1731621.

Barr, Jeremy J., Rita Auro, Mike Furlan, Katrine L. Whiteson, Marcella L. Erb, Joe Pogliano, Aleksandr Stotland, et al. 2013. “Bacteriophage Adhering to Mucus Provide a Non-Host-Derived Immunity.” Proceedings of the National Academy of Sciences of the United States of America 110 (26): 10771–76.

Belkaid, Yasmine, and Timothy W. Hand. 2014. “Role of the Microbiota in Immunity and Inflammation.” Cell 157 (1): 121–41.

Berkson, Julia D., Claire E. Wate, Garrison B. Allen, Alyxandria M. Schubert, Kristin E. Dunbar, Michael P. Coryell, Rosa L. Sava, et al. 2024. “Phage-Specific Immunity Impairs Efficacy of Bacteriophage Targeting Vancomycin Resistant Enterococcus in a Murine Model.” Nature Communications 15 (1): 2993.

Bichet, Marion C., Wai Hoe Chin, William Richards, Yu-Wei Lin, Laura Avellaneda-Franco, Catherine A. Hernandez, Arianna Oddo, et al. 2021. “Bacteriophage Uptake by Mammalian Cell Layers Represents a Potential Sink That May Impact Phage Therapy.” iScience 24 (4): 102287.

Billaud, Maud, Quentin Lamy-Besnier, Julien Lossouarn, Elisabeth Moncaut, Moira B. Dion, Sylvain Moineau, Fatoumata Traoré, et al. 2021. “Analysis of Viromes and Microbiomes from Pig Fecal Samples Reveals That Phages and Prophages Rarely Carry Antibiotic Resistance Genes.” ISME Communications 1 (1): 55.

Billaud, Maud, Marie-Agnès Petit, and Julien Lossouarn. 2023. “The Clostridium-Infecting Filamentous Phage CAK1 Genome Analysis Allows to Define a New Potential Clade of Tubulavirales.” FEMS Microbiology Letters 370 (January): fnad099.

Brzozowska, Ewa, Tomasz Lipiński, Agnieszka Korzeniowska-Kowal, Karolina Filik, Andrzej Górski, and Andrzej Gamian. 2023. “Detection and Level Evaluation of Antibodies Specific to Environmental Bacteriophage I11mO19 and Related Coliphages in Non-Immunized Human Sera.” *Antibiotics (Basel*, Switzerland) 12 (3): 586.

Burckhardt, Juan C., Derrick H. Y. Chong, Nicola Pett, and Carolina Tropini. 2023. “Gut Commensal Enterocloster Species Host Inoviruses That Are Secreted in Vitro and in Vivo.” Microbiome 11 (1): 65.

Chen, Kang, Giuliana Magri, Emilie K. Grasset, and Andrea Cerutti. 2020. “Rethinking Mucosal Antibody Responses: IgM, IgG and IgD Join IgA.” Nature Reviews. Immunology 20 (7): 427–41.

Dąbrowska, Krystyna. 2019. “Phage Therapy: What Factors Shape Phage Pharmacokinetics and Bioavailability? Systematic and Critical Review.” Medicinal Research Reviews 39 (5): 2000–2025.

Dąbrowska, Krystyna, and Stephen T. Abedon. 2019. “Pharmacologically Aware Phage Therapy: Pharmacodynamic and Pharmacokinetic Obstacles to Phage Antibacterial Action in Animal and Human Bodies.” Microbiology and Molecular Biology Reviews 83 (4). 10.1128/MMBR.00012-19.

Dąbrowska, Krystyna, Paulina Miernikiewicz, Agnieszka Piotrowicz, Katarzyna Hodyra, Barbara Owczarek, Dorota Lecion, Zuzanna Kaźmierczak, Andrey Letarov, and Andrzej Górski. 2014. “Immunogenicity Studies of Proteins Forming the T4 Phage Head Surface.” Journal of Virology 88 (21): 12551–57.

Dahlman, Sofia, Laura Avellaneda-Franco, Emily L. Rutten, Emily L. Gulliver, Sean Solari, Michelle Chonwerawong, Ciaren Kett, et al. 2025. “Isolation, Engineering and Ecology of Temperate Phages from the Human Gut.” Nature 647 (8090): 698–705.

Easwaran, Maheswaran, Fatma Abdelrahman, Ayman El-Shibiny, Baskar Venkidasamy, Muthu Thiruvengadam, Periyasamy Sivalingam, Dhanraj Ganapathy, Juhee Ahn, and Hyun Jin Shin. 2025. “Exploring Bacteriophages to Combat Gut Dysbiosis: A Promising New Frontier in Microbiome Therapy.” Microbial Pathogenesis 208 (108008): 108008.

Enault, François, Arnaud Briet, Léa Bouteille, Simon Roux, Matthew B. Sullivan, and Marie-Agnès Petit. 2017. “Phages Rarely Encode Antibiotic Resistance Genes: A Cautionary Tale for Virome Analyses.” The ISME Journal 11 (1): 237–47.

Fitzgerald C. Brian, Andrey N. Shkoporov, Aditya Upadrasta, Ekaterina V. Khokhlova, R. Paul Ross, and Colin Hill. 2021. “Probing the ‘Dark Matter’ of the Human Gut Phageome: Culture Assisted Metagenomics Enables Rapid Discovery and Host-Linking for Novel Bacteriophages.” Frontiers in Cellular and Infection Microbiology 11 (March): 616918.

Gembara, Katarzyna, and Krystyna Dąbrowska. 2021. “Phage-Specific Antibodies.” Current Opinion in Biotechnology 68 (April): 186–92.

Guerin, Emma, and Colin Hill. 2020. “Shining Light on Human Gut Bacteriophages.” Frontiers in Cellular and Infection Microbiology 10 (September): 481.

Haddock, N. L., L. J. Barkal, and P. L. Bollyky. 2023. “Bacteriophage Populations Mirror Those of Bacterial Pathogens at Sites of Infection.” mSystems 8 (4): e0049723.

Hodyra-Stefaniak, Katarzyna, Zuzanna Kaźmierczak, Joanna Majewska, Sanna Sillankorva, Paulina Miernikiewicz, Ryszard Międzybrodzki, Andrzej Górski, et al. 2020. “Natural and Induced Antibodies against Phages in Humans: Induction Kinetics and Immunogenicity for Structural Proteins of PB1-Related Phages.” *PHAGE (New Rochelle*, N.Y.) 1 (2): 91–99.

Kaźmierczak, Zuzanna, Joanna Majewska, Paulina Miernikiewicz, Ryszard Międzybrodzki, Sylwia Nowak, Marek Harhala, Dorota Lecion, et al. 2021. “Immune Response to Therapeutic Staphylococcal Bacteriophages in Mammals: Kinetics of Induction, Immunogenic Structural Proteins, Natural and Induced Antibodies.” Frontiers in Immunology 12 (June): 639570.

Kharrat, Lukeman, Camilo A. Garcia-Botero, Wade Ingersoll, Tiffany Luong, Alejandro Reyes, and Dwayne R. Roach. 2025. “Large-Scale Genomic Analysis of CpG-Mediated Immunogenicity in Bacteriophages and a Novel Predictive Risk Index.” bioRxivorg. 10.1101/2025.05.15.652987.

Knezevic, Petar, Evelien M. Adriaenssens, and Ictv Report Consortium. 2021. “ICTV Virus Taxonomy Profile: Inoviridae.” The Journal of General Virology 102 (7): 001614.

Kuang, Xu, Jamie Gorzynski, Marie Touchon, Andrey Shkoporov, Eduardo P. C. Rocha, J. Ross Fitzgerald, John Chen, Jakob T. Rostøl, and José R. Penadés. 2026. “Bacteriophages Mobilize Bacterial Defense Systems via Lateral Transduction.” Science Advances 12 (4): eadx5749.

Le Huu Thanh, Alicia Fajardo Lubian, Bethany Bowring, David van der Poorten, Jonathan Iredell, Jacob George, Carola Venturini, Golo Ahlenstiel, and Scott Read. 2024. “Using a Human Colonoid-Derived Monolayer to Study Bacteriophage Translocation.” Gut Microbes 16 (1): 2331520.

Liao, Hanpeng, Chen Liu, Shungui Zhou, Chunqin Liu, David J. Eldridge, Chaofan Ai, Steven W. Wilhelm, et al. 2024. “Prophage-Encoded Antibiotic Resistance Genes Are Enriched in Human-Impacted Environments.” Nature Communications 15 (1): 8315.

MacPherson, Chad W., Olivier Mathieu, Julien Tremblay, Julie Champagne, André Nantel, Stéphanie-Anne Girard, and Thomas A. Tompkins. 2018. “Gut Bacterial Microbiota and Its Resistome Rapidly Recover to Basal State Levels after Short-Term Amoxicillin-Clavulanic Acid Treatment in Healthy Adults.” Scientific Reports 8 (1). 10.1038/s41598-018-29229-5.

Majewska, Joanna, Zuzanna Kaźmierczak, Karolina Lahutta, Dorota Lecion, Aleksander Szymczak, Paulina Miernikiewicz, Jarosław Drapała, et al. 2019. “Induction of Phage-Specific Antibodies by Two Therapeutic Staphylococcal Bacteriophages Administered per Os.” Frontiers in Immunology 10 (November): 2607.

Marantos, Anastasios, Namiko Mitarai, and Kim Sneppen. 2022. “From Kill the Winner to Eliminate the Winner in Open Phage-Bacteria Systems.” PLoS Computational Biology 18 (8): e1010400.

Moustafa, Ahmed, Chao Xie, Ewen Kirkness, William Biggs, Emily Wong, Yaron Turpaz, Kenneth Bloom, et al. 2017. “The Blood DNA Virome in 8,000 Humans.” PLoS Pathogens 13 (3): e1006292.

Nguyen, Sophie, Kristi Baker, Benjamin S. Padman, Ruzeen Patwa, Rhys A. Dunstan, Thomas A. Weston, Kyle Schlosser, et al. 2017. “Bacteriophage Transcytosis Provides a Mechanism to Cross Epithelial Cell Layers.” mBio 8 (6). 10.1128/mBio.01874-17.

Niault Theophile, Stineke van Houte, Edze Westra, and Daan C. Swarts. 2025. “Evolution and Ecology of Anti-Defence Systems in Phages and Plasmids.” Current Biology 35 (1): R32–44.

Pfeifer, Eugen, Rémy A. Bonnin, and Eduardo P. C. Rocha. 2022. “Phage-Plasmids Spread Antibiotic Resistance Genes through Infection and Lysogenic Conversion.” mBio 13 (5): e0185122.

Ramírez, Angie L., Luisa Páez, Laura Vega, Viviana Aya, Carolina Hernández, Nicolas Luna, Marina Muñoz, Luz Helena Patiño, and Juan David Ramírez. 2026. “Metagenomic Analysis of the Human Gut Virome Reveals Functional Signatures and Viral Stability across Hospitalized and Non-Hospitalized Diarrheal and Non-Diarrheal Individuals.” Gut Pathogens 18 (1): 17.

Rogier, Eric W., Aubrey L. Frantz, Maria E. C. Bruno, and Charlotte S. Kaetzel. 2014. “Secretory IgA Is Concentrated in the Outer Layer of Colonic Mucus along with Gut Bacteria.” Pathogens 3 (2): 390–403.

Round, June L., and Sarkis K. Mazmanian. 2009. “The Gut Microbiota Shapes Intestinal Immune Responses during Health and Disease.” Nature Reviews. Immunology 9 (5): 313–23.

Rousset, François, Florence Depardieu, Solange Miele, Julien Dowding, Anne-Laure Laval, Erica Lieberman, Daniel Garry, Eduardo P. C. Rocha, Aude Bernheim, and David Bikard. 2022. “Phages and Their Satellites Encode Hotspots of Antiviral Systems.” Cell Host & Microbe 30 (5): 740–753.e5.

Roux, Simon, Mart Krupovic, Rebecca A. Daly, Adair L. Borges, Stephen Nayfach, Frederik Schulz, Allison Sharrar, et al. 2019. “Cryptic Inoviruses Revealed as Pervasive in Bacteria and Archaea across Earth’s Biomes.” Nature Microbiology 4 (11): 1895–1906.

Santiago-Rodriguez, Tasha M., and Emily B. Hollister. 2022. “Unraveling the Viral Dark Matter through Viral Metagenomics.” Frontiers in Immunology 13 (September): 1005107.

Shkoporov, Andrey N., and Colin Hill. 2019. “Bacteriophages of the Human Gut: The ‘Known Unknown’ of the Microbiome.” Cell Host & Microbe 25 (2): 195–209.

Smith, Linda, Ekaterina Goldobina, Bianca Govi, and Andrey N. Shkoporov. 2023. “Bacteriophages of the Order Crassvirales: What Do We Currently Know about This Keystone Component of the Human Gut Virome?” Biomolecules 13 (4): 584.

Sutton, Thomas D. S., and Colin Hill. 2019. “Gut Bacteriophage: Current Understanding and Challenges.” Frontiers in Endocrinology 10 (November): 784.

Zuo, Tao, Xiao-Juan Lu, Yu Zhang, Chun Pan Cheung, Siu Lam, Fen Zhang, Whitney Tang, et al. 2019. “Gut Mucosal Virome Alterations in Ulcerative Colitis.” Gut 68 (7): 1169–79.

Zuppi, Michele, Heather L. Hendrickson, Justin M. O’Sullivan, and Tommi Vatanen. 2021. “Phages in the Gut Ecosystem.” Frontiers in Cellular and Infection Microbiology 11: 822562.

