## Supplementary methods for "Phageome transfer from gut to circulation and its regulation by human immunity"

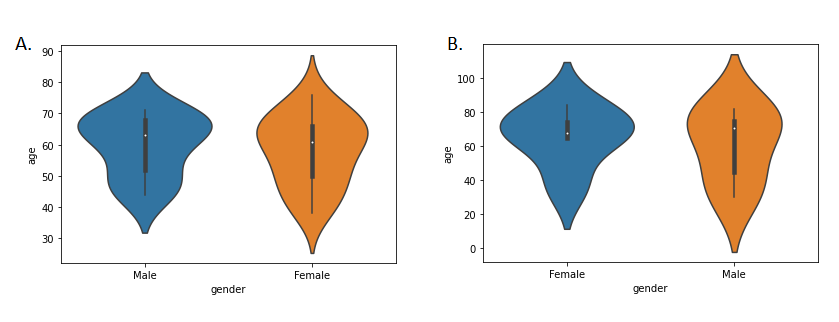

**Figure S1 (A) Gender age profile for the group where at least one case of bacteriophage** transcytosis was identified. The average age for women is 58.1 and for men is 59.9. (B) Gender age profile for the group where at least one case of bacteriophage transcytosis was not identified. The mean age for women is 65.6 and for men is 60.6

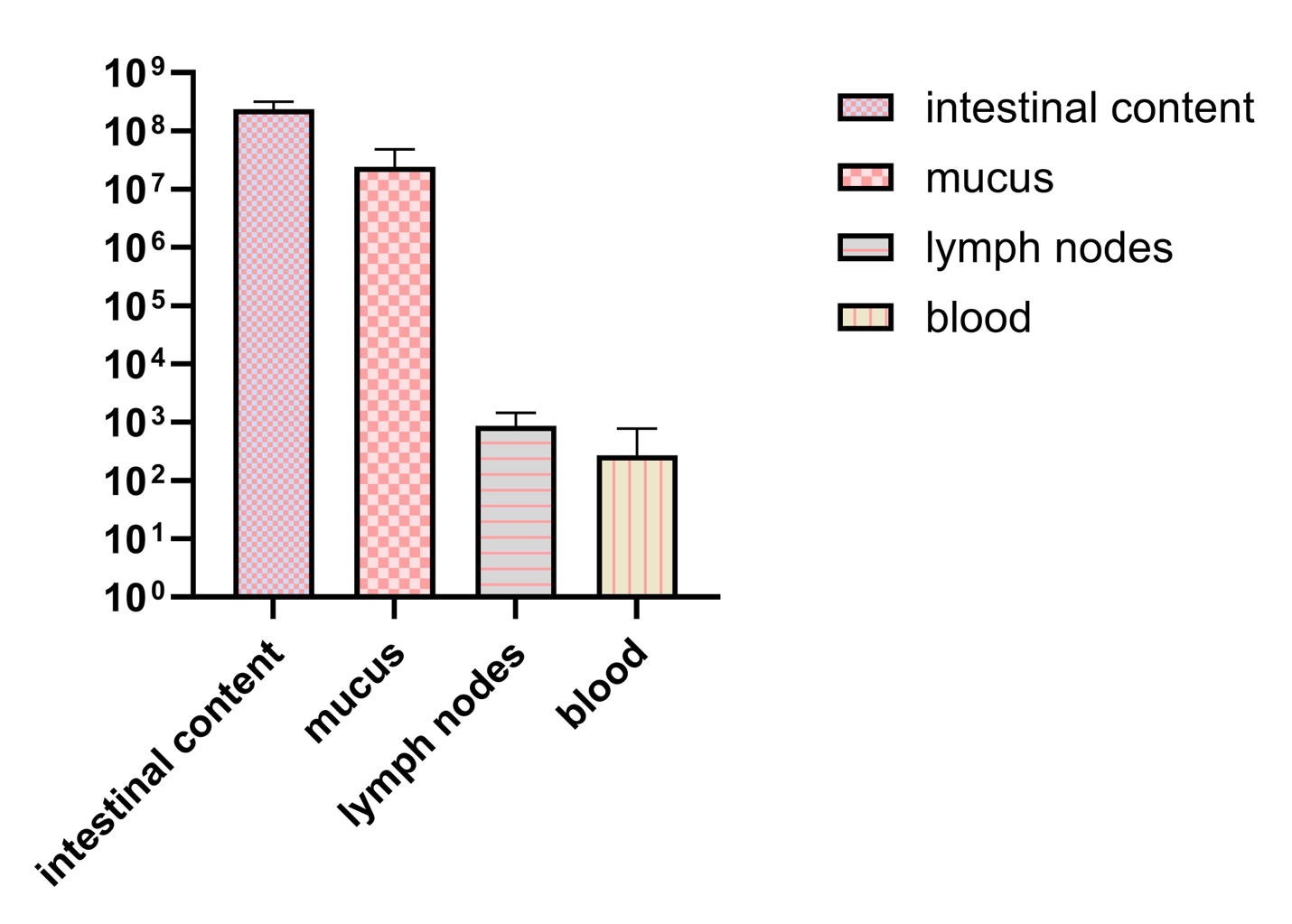

**Figure S2 Binding of T4 phage to the intestinal mucosa of mice followed by phage translocation to blood circulation.** C57 BL/6 female mice (N=6) were given phage T4 preparation purified with Sephacryl in drinking water diluted to a titer of 5x10^9^ pfu/ml; drinking water contained phages during the whole experiment. Blood was sampled from the animals tail vein to heparinized tubes 24 hours after starting phage administration, than the animals were euthanized and intestinal samples were collected: intestinal contents and intestinal mucosal layer (by scrubbing); lymph node tissue was also collected. Phage concentration was assessed using the double-layer agar plate method (100 µl blood per plate); lymphatic tissue or intestinal samples were weighed in PBS (10^-1^ weight-to-volume), homogenized, shaken for 15 mins to release phages, diluted from 10^-1^ to 10^-6^ and quantified using spotting method on bacterial layer.

**Detailed IgG-reactivity – Phagescan protocol**

Identification of bacteriophage oligopeptides interacting with specific IgGs is presented in Figure 1. It was performed in accordance with the modified protocol published by Xu et al., 2015and adapted for our research using coding sequences of the investigated bacteriophages as the source for library design. The GenBank database (National Center for Biotechnology Information) was searched for all available phage genomes (accessed on: June 1, 2018). For further research, annotated open reading frames (ORFs) were selected, which were described with the keywords: 'structural', 'tail', 'decoration', 'capsid', 'scaffold', 'baseplate', 'coat', 'virion', 'structure', ‘lysin’, ‘muramidase’ or ‘holin’ but did not contain the keywords: 'initiator', 'chaperone', 'maturation protease', 'morphogenesis'. Phage enzyme-coding sequences were used as a reference for possible comparisons between structural proteins and propagation cycle-related proteins of phages. The amino acid sequences of selected ORF were virtually cut into 56 aa long fragments tailing through protein sequences, starting every 28 aa from the first amino acid (50% overlap was applied to avoid splitting potential epitopes). Sequences were deduplicated and then reverse-translated into DNA sequences using codons optimized for expression in E. coli; as a result, 277 051 oligopeptide sequences were included in the library. The oligopeptide library was synthesized using the SurePrint technology for nucleotide printing (Agilent). These oligonucleotides were used to create a phage display library using the T7Select 415-1 Cloning Kit (Merck Millipore) using Liga5™ (A&A Biotechnology) instead of T4 ligase and default restriction sites (EcoRI and HindIII).

Specific reactions between phage-derived oligopeptides and antibodies present in the investigated sera were identified by immunoprecipitation. Immunoprecipitation of the library was performed in accordance with a previously published protocol (Xu et al., 2015) . Briefly, the phage library was amplified in a standard culture as described in the manufacturer’s manual and then purified using a hollow fiber filtration membrane. In the final step, the purified and concentrated library was suspended in a Phage Extraction Buffer (20 mM Tris-HCl, 100 mM NaCl, 60 mM MgSO4, pH 8.0). All plastic containers (96-well plates) used for immunoprecipitation were prepared by blocking with 3% Bovine Serum Albumin (BSA) in TBST buffer overnight on a rotator (40 rpm, 4°C). A sample representing an average of 2x10^5^ copies of each clone in 250 µL was mixed with 1 µL of human serum (two technical replicates were applied) and incubated overnight at 4°C with rotation (50 rpm). A 20μL aliquot of a 1:1 mixture of Protein A and Protein G Dynabeads (Invitrogen) was added and incubated for 4h at 4°C with rotation (50 rpm). The liquid in all wells was separated from Dynabeads on a magnetic stand and removed. Beads were washed 5 times with 280µL of a wash buffer (50 mM Tris-HCl, pH 7.5, 150 mM NaCl, 0.1% Tween-20) and beads were resuspended in 60µL of water to elute the immunoprecipitated bacteriophages from the beads.

The immunoprecipitated part of the library was then used for amplification of the insert region according to the manufacturer’s instructions with a Phusion Blood Direct PCR Kit (Thermo Fisher Scientific). The primers T7_Endo_Lib_LONG_FOR GCCCTCTGTGTGAATTCT and T7_Endo_Lib_LONG_REV GTCACCGACACAAGCTTA were used, and a second round of PCR was carried out with the IDT for Illumina UD indexes (Illumina Corp.) to add adapter tags. Sequencing of the amplicons in accordance with Illumina next generation sequencing (NGS) technology was outsourced (Genomed, Warszawa). A FASTA file containing the selected oligonucleotide sequences was utilized to construct a BLAST database. This was achieved by executing the "makeblastdb" command with the "nucl" dbtype option. Reads from sequencing were annotated using BLAST 2.12 algorithm.

**Detailed mulitpanel figures creation detailed descriptions:**

**Figure 3. Taxonomic and host context of vOTUs detected in both intestine and serum.**

**A) Figure Legend**

Fig. 1a–d | **a–b**, Intestinal abundance (log10 scale) of viral operational taxonomic units (vOTUs) stratified by **phage family** (a) or predicted **host class** (b), comparing vOTUs with **Only gut presence** versus **Transcytosed** status (violin distributions with embedded boxplots and overlaid points). Significance within each category was assessed by Wilcoxon tests and displayed as in-panel significance labels with false discovery rate (FDR) correction. **c**, Circular phylogeny of the transcytosed vOTU set with adjacent gapless annotation rings showing **ICTV order** (taxonomy) and predicted **host class**; entries lacking assignment are grouped as **Unknown**. **d**, Counts of transcytosed vOTUs per phage family (stacked by host class), with totals shown above bars and a y-axis break to accommodate the dominant Unclassified bin.

**B) Supplementary Note**

**Panel Purpose**

This multi-part panel links **transcytosis status** to intestinal abundance patterns and taxonomic/host context. Panels **a–b** ask whether intestinal abundance differs between vOTUs detected only in gut versus those also detected in serum (Transcytosed) when stratified by **phage family** or **host class**. Panels **c–d** summarize the phylogenetic breadth of the transcytosed set and quantify how those vOTUs distribute across **phage families** and **host classes**, while retaining unassigned categories under **Unknown**.

**Data and Processing**

Analyses draw from a per-vOTU metadata table (Supplementary Table 1) which contain curated list of transcytosed/translocated vOTU genome IDs
For panels **a–b**, sample abundance columns were parsed by suffix (_intestine, _serum) and reshaped to long format by subject and tissue. A **contig/vOTU-level transcytosis flag** was assigned within subject by summing abundance across intestine and serum: vOTUs with intestine > 0 and serum > 0 were labeled **Transcytosed**, those with intestine > 0 and serum == 0 were labeled **Only gut presence**, and all other cases were excluded from abundance comparisons. Unclassified categories were removed for the abundance stratifications, and categories were retained only if they contained at least one Transcytosed observation.

For panel **c**, a circular tree was generated from the provided Newick phylogeny after filtering tips to the transcytosed list. Tip metadata were joined by genome_id, and two gapless rectangular rings were drawn outside the tree to encode **ICTV order** and **host class**. Missing/empty assignments were consolidated as **Unknown** for display consistency. For panel **d**, the same transcytosed list was used to filter metadata and tabulate vOTU counts per **phage family × host class**, producing stacked bars with per-family totals.

**Design Choices**

Panels **a–b** use violin+boxplots with point overlays to show distribution shape, robust summaries, and per-observation variability on a **log10 y-scale**. Category-wise Wilcoxon tests were computed within each family/host class and multiple-testing controlled by **FDR**, with significance rendered as compact in-panel labels. Panels **c–d** share a single, harmonized encoding of **host class** (including an **Unknown** level), so the stacked composition in **d** can be read directly against the host-class ring in **c**. Panel **d** applies a y-axis break to preserve visibility of low-count families while retaining the dominant Unclassified bin.

**Reading the Panel**

In **a**, compare the Only gut presence versus Transcytosed distributions within each phage family; in **b**, make the same comparison within each host class. Significance markers indicate categories where the two groups differ under the Wilcoxon/FDR framework. In **c**, each tip corresponds to a transcytosed vOTU; the inner ring encodes ICTV-order taxonomy and the outer ring encodes host class, allowing visual assessment of whether specific clades share taxonomy/host annotations or are interspersed. In **d**, bar height gives the total number of transcytosed vOTUs per phage family, and stacked colors partition that total by host class (with Unknown preserved).

**Interpretation**

Together, these panels show that the **transcytosed vOTU set spans multiple phage families and host classes**, and that the set is not confined to a single phylogenetic region of the tree. The combined view supports interpreting transcytosed detections as arising from a **compositionally heterogeneous subset** of gut vOTUs while explicitly retaining uncertainty through the **Unknown** category in both host and taxonomy displays.

**Figure 4 Functional and defense signatures among transcytosed phage contigs.**

**A) Figure Legend**

Fig. 2a–c | Functional and defense-associated features of contigs/protein groups stratified by gut-only detection versus serum co-detection. **a,** Intestinal abundance (log10-transformed) for protein groups, comparing sequences with *Only gut presence* versus *Translocated* status (serum >0 in matched samples). Protein groups are ordered by keyword-assigned structural category (Tail, Capsid, Baseplate, Lysis, Structural), indicated by shaded background blocks and category header bars; significance is assessed per protein group using paired Wilcoxon tests on per-sample medians with Benjamini–Hochberg false discovery rate (FDR) correction, shown as asterisks. **b,** Clustered heatmap of KEGG pathways (ko IDs) by phage family, displaying row-wise Z-scores of summed abundance; rows are annotated by COG functional category and columns by host phylum, with both legends placed below the heatmap. **c,** Bar plots summarizing contigs count for DefenseFinder calls by defense type/activity and AMR gene subclasses.

Panel 2Figure_Panel_A_B_C_publi…

**B) Supplementary Note**

**Panel Purpose**

This panel summarizes three complementary views of functional signals associated with “translocation” as defined in the analysis code: sequences detected in intestines and also detected in serum for the same subject/sample (Translocated) versus sequences detected only in intestines (Only gut presence). Panel A tests whether intestinal abundance differs between these two detection states across protein groups while organizing protein groups into structural/functional categories. Panel B contextualizes functional annotation patterns by relating KEGG pathways to phage families, with row and column metadata highlighting broad COG categories and host phyla. Panel C provides a compact overview of defense/anti-defense system calls and antimicrobial resistance (AMR) subclass annotations present in the integrated dataset.

**Data and Processing**

For **Panel A**, a long-format table is constructed from merged_all_scan_names by identifying paired serum and intestine columns using suffixes (_serum, _intestine) and intersecting sample identifiers to define matched sets. For each protein group and shared sample, the intestinal value is retained when >0 and serum is used to classify status (serum >0 → Translocated; serum 0/NA → Only gut presence). Intestinal values are log10-transformed as log10(intestine_value + 1), and only protein groups that exhibit at least one translocated observation are retained. Protein groups are assigned to one of five categories (Tail, Capsid, Baseplate, Lysis, Structural) using a keyword-based classifier; uncategorized entries are dropped (no “Other” class). Significance is computed per protein group using paired Wilcoxon tests on per-sample median log10 intestinal values (Translocated vs Only gut presence), followed by Benjamini–Hochberg FDR correction; adjusted p-values are converted to star annotations.

For **Panel B**, Supplementary Table 2 is filtered to retain rows with non-zero abundance across sample columns and with KEGG pathway strings containing valid koXXXXX identifiers. Rows are exploded to one KEGG term per record, and a KEGG × phage family matrix is created using summed total abundance across all serum and intestine sample columns. The heatmap displays row-wise Z-scores (with a safeguard returning zeros for constant rows) and is hierarchically clustered on both rows and columns. Row color annotations are assigned by the modal COG category per KEGG pathway (mapped to full COG category names), and column color annotations are assigned by the modal host phylum per phage family. The x-axis label is “Phage family”, the figure title is suppressed, and both legends are placed below the heatmap for readability.

For **Panel C**, defense counts are computed from defense_csv as the number of records per (type, activity), and AMR subclass counts are computed from the Subclass field in Supplementary Table 2 Both bar plots annotate counts above bars; the y-axis label is set to “Contigs count”.

**Design Choices**

Panel A uses log10(intestine+1) to stabilize scale and allow inclusion of small positive values, and it reduces within-sample variability by testing paired differences on per-sample medians before applying a paired Wilcoxon test. Multiple testing is handled using Benjamini–Hochberg FDR, and results are communicated with conventional star thresholds. The categorical organization in Panel A is intentionally rule-based (keyword matching) to enforce consistent grouping and to remove ambiguous “Other” assignments.

Panel B uses row-wise Z-scoring to emphasize relative pathway patterns across phage families rather than absolute abundance, which is already aggregated across all serum and intestine sample columns. Legends are positioned below the heatmap (and the “COG category” label is kept horizontal in the final layout) to prevent overlap with tick labels and to preserve heatmap width.

**Reading the Panel**

In **Panel A**, each protein group has two distributions (Only gut presence vs Translocated) shown as boxplots with overlaid jittered points; category blocks and header bars indicate where Tail, Capsid, Baseplate, Lysis, and Structural groups fall along the x-axis. Asterisks beneath protein groups indicate FDR-adjusted significance of paired differences. The accompanying horizontal alluvial-style summary (within Panel A) reports category totals for all categorized contigs versus translocated contigs and visualizes category-wise flow between these totals.

In **Panel B**, the heatmap rows (KEGG pathways) and columns (phage families) are clustered; warmer/cooler colors reflect positive/negative Z-scores within each pathway row. The left-side row color strip encodes COG category, while the top color strip encodes host phylum; both legend keys appear below the heatmap.

In **Panel C**, the upper bar plot separates defense system calls by type and activity (Defense vs Antidefense), while the lower bar plot lists AMR subclasses; bar-top labels are raw counts and the y-axis is “Contigs count”.

**Interpretation**

Together, these subpanels provide a structured, publication-ready summary of how (i) intestinal abundance by protein group relates to serum co-detection under a paired-sample framework (Panel A), (ii) functional pathway signatures distribute across phage families while tracking broad functional and host metadata (Panel B), and (iii) defense/anti-defense and AMR subclass annotations are represented in the integrated callsets (Panel C). Key constraints are embedded in the implementation: “translocation” is defined strictly by non-zero detection in matched serum and intestine columns (without an explicit abundance threshold beyond >0), functional categories in Panel A depend on keyword matching, and Panel B aggregates abundance across all available sample columns prior to Z-scoring, so it is best read as a pattern map rather than a per-subject effect estimate.

**Figure 5. IgG-reactivity across gut–serum phage proteins and association with translocation**.

**A) Figure Legend**

Fig. 1a–d | Integrated view of phage protein detection in intestine, serum and IgG-reactivity signals. **a,** Venn diagram of protein-group presence called as values >0 in intestinal metagenomes, serum metagenomes and IgG-reactivity (raw-count signal), highlighting the intestine∩IgG-reactivity-only subset (n=6; 62 intestine-only, 102 IgG-reactivity-only, 69 intestine–serum, 26 shared across all three and 6 intestine∩IgG-reactivity only, as plotted). **b,** Protein-group summary scatterplot showing mean intestinal abundance (x) versus mean serum abundance (y); point colour encodes mean IgG-reactivity (raw counts) and point size reflects log10(total IgG-reactivity+1). **c,** Dual clustered heatmap of log1p-transformed IgG-reactivity reads, row z-scored within each protein label, across predicted lifecycle classes and phage families. **d,** Association of IgG-reactivity (raw reads per subject–category) with translocation score log2((intestine+1)/(serum+1)); regression line shown with Spearman’s ρ and mixed-effects model β (P values) annotated for stratifications by phage host, phage cycle and phage protein function. IgG, immunoglobulin G.

**B) Supplementary Note**

**Panel Purpose**

This multipanel figure links three layers of evidence—compartmental detection (intestine versus serum), antibody-associated signal (IgG-reactivity) and abundance-based contrast between compartments (translocation score)—to contextualize which phage protein groups are detected where, which are IgG-reactive, and how IgG-reactivity covaries with intestine-to-serum abundance differences when summarized by biological groupings (host, lifecycle and protein function labels).

**Data and Processing**

Presence/absence in **panel a** is computed directly from a shared table indexed by protein identifiers, using three numeric columns: original_intestine, phagescan_data and original_serum. Values are coerced to numeric and a protein is marked present if it exceeds a threshold of 0.0 in the corresponding column. Sets are formed from the index and intersected to quantify overlaps; the intestine∩IgG-reactivity-only subset (intestine yes, IgG-reactivity yes, serum no) is also exported to a CSV for downstream inspection.

For **panel b**, protein-level observations are first aggregated to protein_group. Intestinal and serum abundance are derived from all columns ending in _intestine and _serum, respectively, with missing values replaced by 0 prior to averaging. IgG-reactivity is aggregated from PhageScan-style donor columns identified as purely numeric column names; mean IgG-reactivity is computed as the mean of raw counts across donors, and total IgG-reactivity is the sum across donors. A log10(total+1) transform is used for point-size scaling.

**Panel c** visualizes two matrices (sharing the same row index of protein-function labels) assembled for lifecycle and family categories, respectively. Values are log1p-transformed IgG-reactivity reads, then standardized as row z-scores. Row and column dendrograms are computed via hierarchical clustering and rendered above/left of each heatmap, with a shared colour scale centred at zero.

For **panel d**, counts are grouped by one of three category columns (phage_host_norm2, phage_ _lifecycle, protein_label) and summed across matched intestine/serum sample columns per subject. Only subject–category pairs with positive values in both compartments (min_value 0.0) are retained, and the translocation score is computed as log2((intestine+1)/(serum+1)). IgG-reactivity is computed as summed raw reads per subject–category from numeric donor columns and filtered to immuno_count >1.0. Associations are summarized using Spearman correlation and a mixed-effects model with subject as the grouping factor (with an optional category variance component).

**Design Choices**

A zero threshold for presence (panel a) keeps the presence call consistent with the boolean logic used elsewhere in the pipeline and ensures that any non-zero signal contributes to overlap structure. Replacing missing abundance values with 0 before averaging (panel b) avoids discarding sparse protein groups while keeping the scale interpretable as a mean abundance across available samples. The combination of mean IgG-reactivity (colour) and total IgG-reactivity (size) separates “broadly reactive” from “strongly reactive” groups without conflating these summaries.

Row z-scoring of log1p IgG-reactivity (panel c) focuses interpretation on *relative* enrichment across categories within each protein label, rather than absolute magnitude differences between labels. In panel d, the +1 pseudocount stabilizes log-ratios when one compartment is low and matches the implemented computation, while the subject-level grouping in the mixed model acknowledges repeated observations within individuals.

**Reading the Panel**

Start with **panel a** to locate which protein groups are detected across data types and to identify the specific intestine∩IgG-reactivity-only subset (n=6) highlighted by the overlap region. Use **panel b** to interpret how protein groups distribute in the intestine–serum abundance space: points near the diagonal reflect similar mean abundance in both compartments, while points displaced toward the x- or y-axis indicate intestine- or serum-weighted means; colour and size reveal which groups carry stronger IgG-reactivity signal.

In **panel c**, read across each row (protein label) to see which lifecycle classes or families are relatively enriched for IgG-reactivity signal after within-row standardization; dendrograms group labels and categories with similar patterns. Finally, **panel d** evaluates whether higher IgG-reactivity at the subject–category level tracks with higher or lower translocation scores; each panel provides both a non-parametric association (ρ) and a model-based slope (β) with P values.

**Interpretation**

Together, these panels provide a structured comparison of detection overlap, abundance contrasts and IgG-reactivity summaries at multiple aggregation levels. Panel d indicates weak or modest associations as plotted (by host: ρ = −0.26, P = 9.34e−02; by cycle: ρ = −0.09, P = 5.69e−01; by protein function: ρ = 0.14, P = 2.03e−01), while the mixed-effects slope term is reported within each subplot (including a significant P value in the lifecycle stratification as annotated). Because panel c uses row z-scores, it should be interpreted as relative enrichment patterns rather than absolute IgG-reactivity magnitudes; similarly, the presence calls in panel a reflect thresholded detection (>0) and do not encode quantitative abundance.

**Figure 6. Co-transcytosis association network and sequence signatures of transcytosed viral contigs**

**A) Figure Legend**

Fig. 1a–d | Multi-panel summary of sequence signatures and co-enrichment structure of intestinal phage material detected in serum. **a,** Unknown–known co-enrichment network where nodes are viral contigs/vOTUs; “Known” nodes are contigs with ICTV assignment at confidence ≥70, and “Sequencing dark matter” are contigs lacking reliable ICTV assignment (family missing/unassigned and/or confidence <70). Edges denote subject-level co-enrichment in serum defined as serum abundance ≥1 and log2((serum+0.5)/(intestine+0.5)) ≥1, retained if co-occurred in ≥3 subjects, φ ≥0.25, and BH-FDR q ≤0.05, with up to 3 edges per unknown. Node size scales with degree; edge width scales with φ; labels shown only for top-degree known nodes. **b,** CpG residuals (GC+log10 length–adjusted log(CpG O/E)) versus log10(contig length, bp), colored by “Transcytosed” vs “Intestines only”. **c,** Group shifts in sequence features (log10 length, GC proportion, mono-nucleotide Shannon entropy, ORF density) with Wilcoxon rank-sum p-values and BH-FDR. **d,** Known families ranked by number of unique unknown partners (q ≤0.05), with edge counts annotated.

**B) Supplementary Note**

**Panel Purpose**

This panel tests whether viral contigs detected in serum (“transcytosed”) exhibit distinct sequence composition relative to contigs detected only in intestines, and whether unclassified viral sequences (“sequencing dark matter”) show structured co-enrichment relationships with taxonomically assigned phage families across subjects.

**Data and Processing**

Inputs comprise a vOTU representative FASTA (vOTU_reps.fasta), a list of translocated vOTUs (translocated_vOTUs_genomes.txt), and a tidy per-subject table containing paired intestine and serum abundances with ICTV taxonomy fields (translocated_vOTUs_taxonomy_tidy.tsv). Sequence-derived features were computed directly from nucleotide sequences: length (bp) and log10(length), GC proportion, mono-nucleotide Shannon entropy (bits) from A/C/G/T frequencies, and a simple ORF density estimate on the plus strand based on ATG start and TAA/TAG/TGA stop codons with a minimum ORF length of 30 aa. CpG observed/expected (CpG O/E) was computed from observed “CG” dinucleotide counts relative to the expectation from p(C)×p(G)×(L−1), then transformed as log(CpG O/E + ε) with ε=1e−6. CpG residuals were obtained by fitting a linear model log(CpG O/E + ε) ~ GC + log10(length) and taking per-contig residuals. Group membership for panels b–c was defined by the translocated ID list, displayed as “Transcytosed” versus “Intestines only”.

**Design Choices**

Unknown sequences were defined using ICTV annotation fields as contigs with missing/unassigned family and/or ICTV confidence <70; known sequences required ICTV confidence ≥70. Subject-level serum enrichment was defined to avoid ratio inflation at low counts using thresholds serum ≥1 and log2((serum+0.5)/(intestine+0.5)) ≥1. Unknown–known associations were quantified across subjects using co-occurrence of enrichment calls, retaining edges with co-occurrence in ≥3 subjects, φ ≥0.25, and BH-FDR q ≤0.05. To prevent visually dense hubs driven by many weak links, edges were capped at 3 per unknown using strongest associations (sorted by q then φ), and unknown nodes were additionally filtered by minimum enriched prevalence (≥0.10 across subjects). Nonparametric group comparisons in panel c used Wilcoxon rank-sum tests with BH-FDR across the four features. For visualization of CpG residuals, the y-axis was clipped to a high quantile of |residual| to improve readability while preserving the underlying residual computation.

**Reading the Panel**

Panel a shows the co-enrichment network with node color indicating taxonomy status (Known vs Sequencing dark matter). Edge thickness reflects association strength (φ), and node size reflects degree (number of retained partners). Only known nodes are labeled, using their identified taxonomic name, limited to the top-degree nodes to maintain legibility; the label count is user-configurable at runtime. Panel b plots CpG residuals against log10 length; points are colored by translocation status, enabling assessment of whether transcytosed contigs systematically deviate in CpG beyond GC/length expectations. Panel c summarizes direction and magnitude of feature shifts: boxplots with overlaid points compare distributions between groups, and the corresponding p-values and BH-FDR are placed at the right edge of each subplot with vertical orientation. Panel d ranks known families by the number of unique sequencing-dark-matter partners among significant edges, with the total number of edges per family annotated for context.

**Interpretation**

This figure jointly supports two inferences. First, transcytosed contigs can be evaluated for sequence-composition signatures independent of trivial confounders (length and GC) by using CpG residuals rather than raw CpG O/E, which is strongly influenced by base composition and genome size. Second, dark-matter contigs do not appear as isolated detections; instead, they form statistically enriched co-enrichment links with known taxa under a per-subject serum-enrichment definition, suggesting non-random co-translocation patterns. The network is constrained by explicit thresholds (serum minimum, log2 fold-change, co-occurrence, φ, and FDR) and an edge-per-unknown cap that favors interpretability over completeness. As implemented, the network encodes association, not causality: shared co-enrichment could reflect shared exposure, shared detection biases, or biological coupling, and the strength metric (φ) is descriptive of co-occurrence across subjects under the chosen enrichment calls.
