## Supplementary QC for "Phageome transfer from gut to circulation and its regulation by human immunity"

### Supplementary QC: Sequencing depth vs translocated vOTUs

#### Key counts

- Per-library depth rows: 74
- Subjects with depth-by-compartment data: 37
- Subjects with translocated vOTU counts: 37
- Subjects in depth-vs-translocated merge: 37

#### Depth vs translocated vOTUs

- Pearson r = 0.205, P = 0.224
- Spearman rho = 0.070, P = 0.68

#### Definition

- Paired-subject depth = sum of post-host-depletion read pairs across intestine and serum (R1).
- Translocated vOTUs counted as (serum > 0.0) AND (intestine > 0.0) within subject.

#### parsing_report.txt

```
Parsed 74/74 R1 files successfully.
```

#### n_report.txt

```
R1 files total: 74
R1 files parsed: 74
Subjects with any depth data: 37
Subjects with BOTH compartments for paired depth plot: 37
Subjects in depth-vs-translocated correlation (inner-join): 37
```

Sequencing depth distributions by compartment (post-host-depletion read pairs)

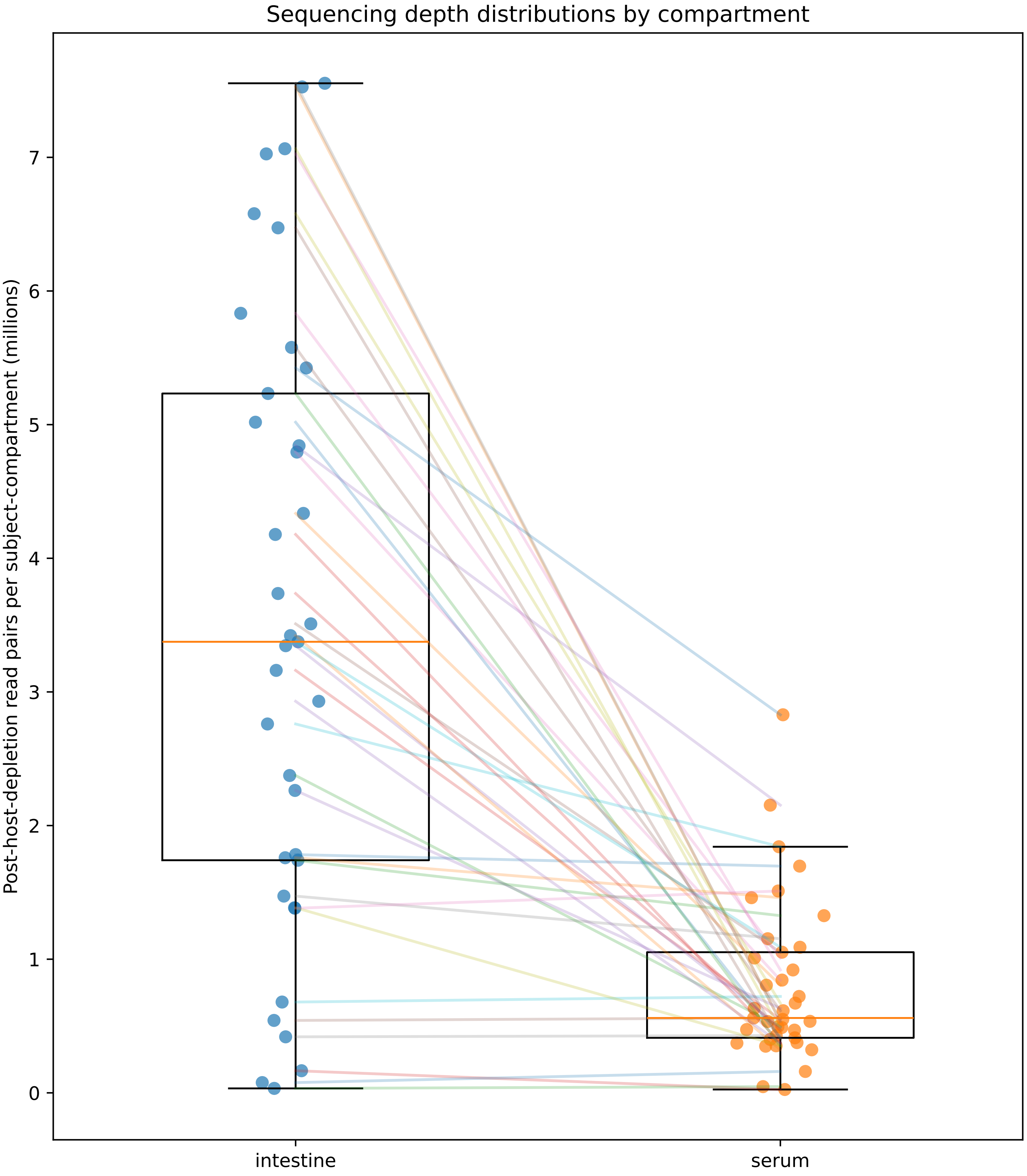

### Sequencing depth vs translocated vOTU detection

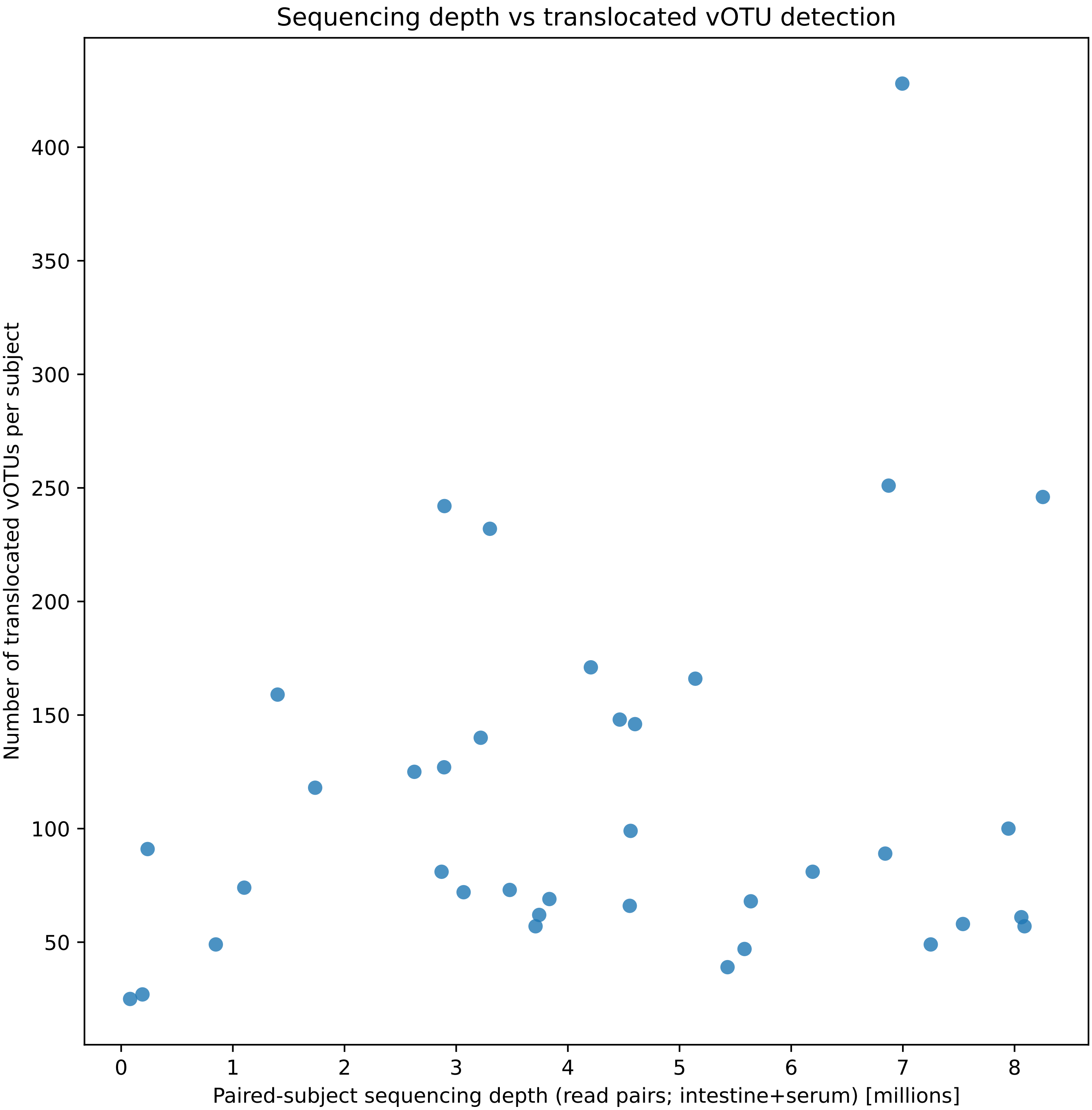


Pearson r=0.205, P=0.224; Spearman rho=0.070, P=0.68. n=37 subjects. Translocated vOTUs counted as (
0.0) AND (intestine > 0.0) within subject.

#### Summary statistics

Depth by compartment (read pairs per subject-compartment)

| compartmer | count | median | mean | min | max |
|---|---|---|---|---|---|
| intestine | 37 | 3,375,155 | 3,507,381 | 33,300 | 7,554,142 |
| serum | 37 | 560,149 | 802,082 | 25,111 | 2,828,980 |

Full per-library depth table is provided as: supp_table_sequencing_depth_post_host_depletion.tsv (recommended to deposit as Supplementary Data / Source Data; not embedded here due to length).
